# Selective Pharmacological Blockade of GPR39 Markedly Reduces No Reflow and Infarct Volumes in a Rat Acute Myocardial Infarction When Administered Prior to Reperfusion

**DOI:** 10.64898/2026.01.16.699339

**Authors:** Carmen Methner, Mary Plascencia, Allura Thompson, Lijuan Liu, Masaki Kajimoto, D. Elizabeth Le, Agostino Cianciulli, Annalisa Pellacani, Jessica Franchi, Chiara Marcheselli, Alberto Parazzoli, Fabrizio Micheli, Sanjiv Kaul

**Author notes:** Presented in part at the Annual Scientific Sessions of the American Heart Association, November 2023 in Philadelphia, and November 2025 in New Orleans, USA. Corresponding Author: Sanjiv Kaul, MD, Knight Cardiovascular Institute Oregon Health and Science University UHN-62, 3181 SW Sam Jackson Park Rd Portland, Oregon 97239.

## Abstract

Our aim was to determine whether selective pharmacological blockade of GPR39 by the novel drug, VC108, reduces no reflow (NRV) and infarct (INV) volumes during acute myocardial infarction (AMI). Immuocytochemistry and qPCR of isolated rat cardiac cells as well as immunohistochemistry and western blot of rat myocardium was performed for presence of GPR39. Rats underwent 1 h of coronary occlusion and 1 h of reperfusion. Groups 1 and 2 animals received drug/vehicle prior to or during coronary occlusion. Groups 3 and 4 received drug/vehicle 5 min prior to or 30 min after reperfusion. Readouts also included tissue pO_2_, hemodynamics, and wall thickening. In Groups 5 and 6 animals, drug was injected for measurement of plasma and tissue levels. Immunocytochemistry and qPCR of cells and immunohistochemistry and western blot of tissue revealed GPR39 expression in all cardiac cells analyzed as well as entire myocardial tissue. There was marked reduction in NRV and INV in groups 1 and 3 animals where both were measured and in Group 2 where INV was measured. In contrast, Group 4 animals failed to show reduction in NRV and INV with the drug. The reduction in NRV in all animals was associated with higher tissue pO_2_ in VC108 compared to vehicle treated animals. Similar results were obtained for INV in only in Group 2 animals. In Group 3 animals direct cardiomyocyte effect of VC108 was seen in myocardium as evidenced by reduced necrosis and apoptosis. We conclude that VC108 is very effective in reducing INV and NRV in an AMI model when given before coronary occlusion or just prior to reperfusion (the latter being clinically more relevant) both in male and female rats. This effect is not seen after reperfusion. VC108 acts by blocking GPR39, resulting in vasodilation through pericyte and VSMC relaxation. It also directly protects cardiomyocytes by preventing downstream effects of GPR39 stimulation.

**New and Noteworthy:** GPR39 is the receptor for 15-HETE, which is a vasoconstrictor with direct cardiomyocyte detrimental effects. Pharmacological inhibition of GPR39 by a novel inhibitor, VC108, reduces coronary no reflow after acute myocardial infarction by relaxing contracted pericytes surrounding capillaries, whereby increasing oxygen delivery. GPR39 inhibition also reduces necrosis and ferroptosis by interrupting aberrant downstream signaling responsible for cell death. Hence, pharmacological inhibition of GPR39 by VC108 offers a novel treatment of acute myocardial infarction.

**Graphical Abstract:** Proposed mechanism of action of VC108 when given prior to and during coronary occlusion based on our results. The drug inhibits the action of the vasoconstrictor, 15-HETE, on GPR39. This causes vasodilation by relaxing contracted pericytes and increasing capillary perfusion, resulting in increased tissue pO_2_ and reduction in no reflow (left side). When tissue pO_2_ is not associated with necrosis, GPR39 inhibition by VC108 directly affects cardiomyocytes by inhibiting downstream signaling of GPR39 present in cardiomyocytes that can lead to cell injury and death. Hence, less necrosis and apoptosis are noted in VC108 versus vehicle treated animals. Created in https://BioRender.com.

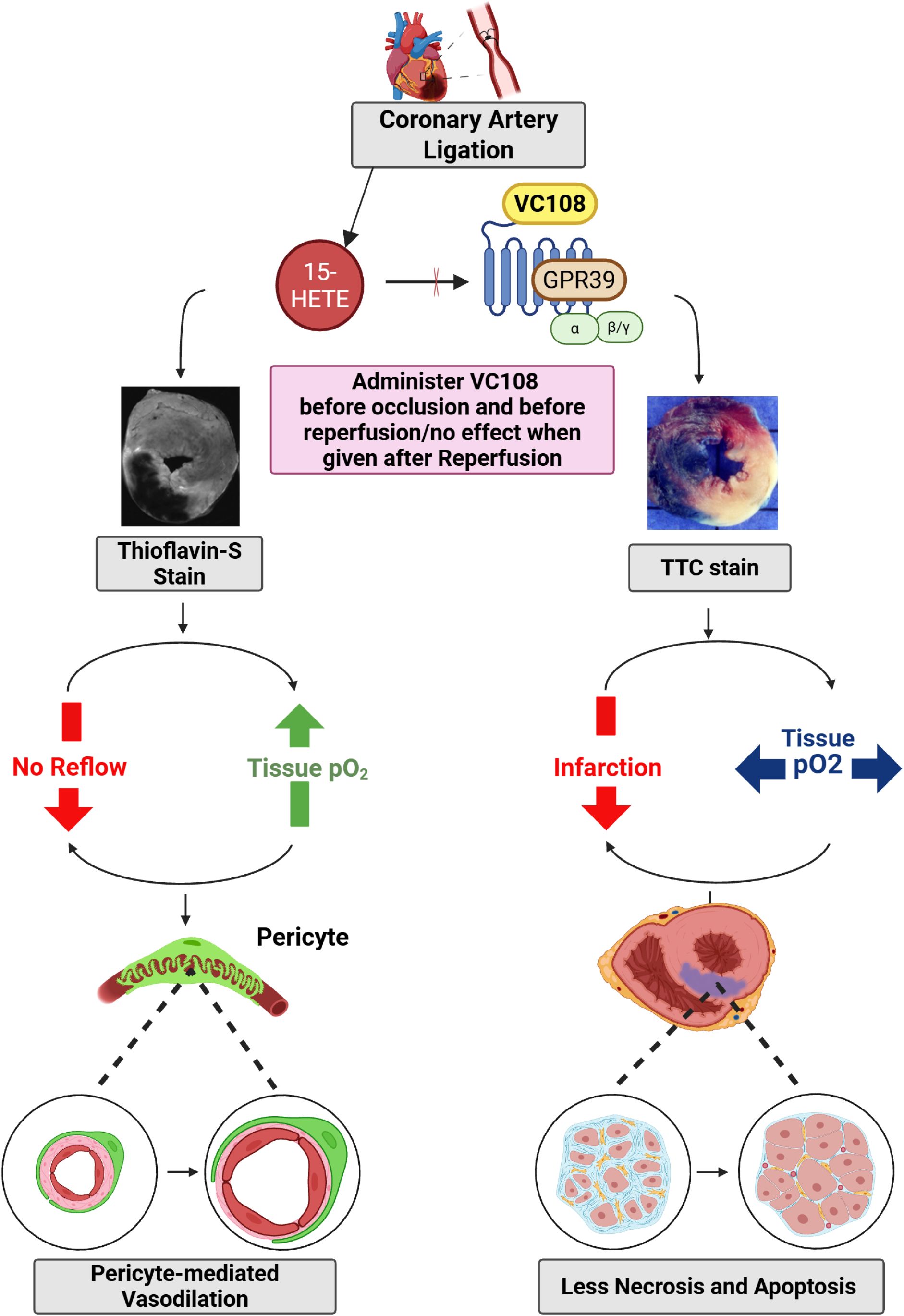

## Introduction

We previously reported that the G-protein coupled receptor 39 (GPR39) is present in the arteriolar vascular smooth muscle cells (VSMCs) and pericytes of the mouse heart where it is the target of the eicosanoids: 15-hydroxyeicosatetraenoic acid (15-HETE), a vasoconstrictor, and 14,15-epoxyeico-satrienoic acid (14,15-EET), a vasodilator.^1^ 15-HETE is the endogenous agonist for GPR39 and causes VSMC and pericyte contraction by increasing cytosolic Ca^++^, an effect that is abolished by pre-treatment with 14,15-EET.^1,2^ GPR39 knock-out mice develop smaller infarct volumes (INV) and no reflow volumes (NRV) compared to wild-type mice subjected to acute myocardial infarction (AMI).^2^ Capillary density and diameter within the ischemic risk area are greater in GPR39 knock-out mice compared to wild-type mice.^2^

GPR39 exhibits significant constitutive activity via its Gα_q_ and Gα_12/13_ subunits.^3,4^ GPR39 activation by synthetic analogs and high doses of Zn^++^ also stimulates Gα_s_ causing cAMP response element-binding protein transcription induction^5,6^ leading to altered expression of genes associated with apoptosis and cell death.^7^ GPR39 activation also stimulates Gα_12/13_, which induces serum response element-mediated transcription that affects cell survival.^8^ Activated Gα_12/13_ stimulates the AKT-phosphoinositide 3-kinase pathway, thus affecting nuclear transcriptional factors causing cell growth and proliferation as well as reducing apoptosis.^9,10^ Additionally, Gα_12/13_ activation through the Rho kinase pathway causes cell proliferation and growth.^8^ 15-HETE activates GPR39 at nanomolar concentration in the presence of physiological concentrations of Zn^++^^1,2^ that acts as a positive allosteric modulator.^3,4^ In our previous experiments, Gα_q_ stimulation, in addition to mobilizing intracellular Ca^++^ causes activation of mitogen-activated protein kinase (MAPK)-extracellular signal-related pathway^1^, increasing cell growth and proliferation.^8^ Genetic deletion of GPR39 resulted in partial or complete inactivation of these pathways in a mouse model of pulmonary hypertension.^11^

We now have a drug candidate that has successfully completed pre-clinical studies for investigational new drug approval for administration as a single intravenous bolus for the treatment for AMI. It is a selective GPR39 antagonist with favorable medicinal chemical properties and has sufficient solubility, and good pharmokinetic properties for target occupancy.^12^ After testing against a panel of putative cardiovascular receptors, including those involved in regulation of vascular tone, the compound was found to be selective for GPR39.^13^ VC108 increases resting coronary blood flow without affecting systemic or pulmonary hemodynamics in the normal dog.^13^

Consequently, we tested the hypothesis that by inhibiting pericyte and VSMC contraction, VC108 will increase tissue O_2_ levels by enhancing tissue perfusion and thus reducing no-reflow volume (NRV) and infarct volume (INV) in a rat model of AMI. To ascertain whether some of the benefits of GPR39 inhibition could result from the presence of the receptor in cardiomyocytes and to establish the presence of GPR39 in rat VSMCs and pericytes we performed immunocytochemistry (ICC) and quantitative polymerase chain reaction (qPCR) of isolated rat cardiomyocytes, pericytes and VSMCs as well as immunohistochemistry (IHC) and Western Blot (WB) of the myocardium for the presence of GPR39. To detect any direct effects of VC108 on cardiomyocytes, we assessed for reduction in necrosis, apoptosis, and ferroptosis compared to vehicle in a subset of animals subjected to AMI. We also performed mass spectroscopy to measure drug levels in plasma during different time points after coronary occlusion or reperfusion, and in ischemic as well as non-ischemic myocardium.

## Materials and Methods

All animal procedures were approved by the Institutional Animal Care and Use Committee of the Oregon Health & Science University and adhered to NIH Guidelines for the Care and Use of Laboratory Animals. All reporting is based on ARRIVE guidelines. Three hundred and fifty-seven wild-type Sprague Dawley rats (8-12 weeks old) of both sexes were used for the in-vitro and in-vivo studies. They were obtained from Charles River Laboratories (Reno, NV, USA). Additionally, 63 Sprague Dawley rats were used for pharmacokinetic studies performed at Aptuit, Verona, Italy.

### In Vitro Experiments

Twelve animals (6 males, 6 females) were used for in vitro experiments, that included ICC of isolated cardiac cells as well as IHC, qPCR and WB of heart tissue. Before removing hearts from the animals, they were anesthetized with 3% isoflurane and euthanized by cervical dislocation.

#### Cell Isolation and culture

Cardiomyocytes were isolated using the Langendorff perfusion apparatus.^14^ They were anticoagulated by intraperitoneal injection of heparin (300 IU) for 20 min before being anesthetized with 3% isoflurane. They were then euthanized using cervical dislocation. The hearts were arrested with cold cardioplegic solution (in mmol/L: 110 NaCl, 16 KCl, 16 MgCl2, 10 NaHCO3, 1.5 CaCl2) administered into the LV lumen after opening the chest cavity from below the rib cage. The hearts were immediately removed from the chest cavity, and the aorta was cannulated with a 14-g blunt-ended under a dissection microscope. The total time for surgery and aortic cannulation was <4 min. The canulated heart was then connected to a pressure constant Langendorff perfusion apparatus that includes a heating coil and a heated circulating bath. The heart was retrogradely perfused with calcium-free perfusion buffer (in mmol/L: 135 NaCl, 4 KCl, 1 MgCl2, 0.33 NaH2PO4, 10 HEPES, 5 Taurine, 20 BDM (2,3-butanedione monoxime), and 10 Glucose, pH adjusted to 7.4 with NaOH) for 3 min at a constant pressure of 70 mmHg. Subsequently, the perfusate was switched to digestion buffer with 0.3 mg/mL collagenase B (COLLB-RO, Sigma, St. Louis, MO, USA), 0.4 mg/mL collagenase D (COLLD-RO, Sigma), and 0.05 mg/mL protease XIV (P5147, Sigma) at a temperature of 37°C for 30 min. Afterward, the atria and large vessels were removed, and the cardiac ventricles were cut into small pieces and further digested 3 times for 5 min each at 37°C using 0.05% Trypsin-EDTA (Thermo Fisher scientific) in a small beaker under gentle agitation.

After centrifuging at 20 G for 2 min, tissue was collected in a petri dish containing the perfusion buffer with 1% bovine serum albumin (A4503, Sigma). Cardiomyocytes were filtered through a 200 µm filter to remove any undigested tissue and then enriched by gravity for 15 min and followed by the gradual addition of CaCl2 (final concentration of Ca^++^ was 1 mM). The cells were plated on uncoated dish with culture medium (DMEM, 1% penicillin-streptomycin, and 10% FBS) and incubated at 37°C and 5% CO2 for 1 h to remove non-cardiomyocytes. The isolated cardiomyocytes (non-adhesive cells) were plated on 25 mm coverslips pre-coated with 10 μg/mL laminin (L2020, Sigma) in 6 well plates for 2 h.

For VSMC isolation, after removing the vessels, hearts were placed into a coronal heart slicer matrix, where 4 cross-sectional slices were made from each heart.^15^ The slices were placed into a collagen (Sigma-Aldrich, St. Louis MO, #C5533)-coated tissue culture plate, 1 slice per well of a 24-well plate, containing 100 µL FBS. The culture plate was placed into a cell culture incubator (37 °C; 95% air and 5% CO2) for 4 h after which, 500 µL of VSMC culture medium was added to each well. The medium was changed 5 days later and incubated for a further 7 days, allowing for migration of VSMCs from the heart slices onto the tissue culture plastic. The slices were carefully removed from the wells using forceps (Dumont #5) and discarded. Cells were enzymatically detached from wells using trypsin: EDTA (0.05%: 0.5 M), once cells had detached 0.2 mL DMEM+10% FBS per well was used to inactivate the enzyme. The cell suspension was transferred to a 15 mL conical tube, pooled, and centrifuged at 1000 rpm for 8 min. The supernatant was removed, cells were re-suspended in 10 mL of SMC culture medium.

For pericyte isolation, ventricles were diced and digested with collagenase (CLS-2, Worthington, Cat. #LS004176) in an agitated water bath (37°, 100 RPM) for 70 min.^15^ This mixture was triturated using a 14-gauge metal cannula attached to a 35 mL disposable syringe every 20 mins. After the final trituration, the suspension was passed through a 70 µm disposable cell strainer and collected in a 50 mL conical tube. Additionally, 10 mL DMEM was passed through the cell strainer to collect any remaining cells. The cell suspension was centrifuged at 1000 rpm for 10 min at room temperature. To isolate pericytes, the cells were resuspended in 6 mL DMEM + 60 µL CD31 (BD Pharmingen, Cat. #553369) conjugated Dynabeads (Sheep anti-Rat IgG, Cat. #11035) in a 15 mL conical tube and placed on a rotator (medium setting) for 40 min at RT. The tube was mounted into a magnetic separator for 1 min allowing the beads (with cells attached) to adhere to the sides. The supernatant containing unbound cells (CD31-) was collected and further isolated with PDGFRβ CD140b (Invitrogen, Cat. #14-1402-82) using the same method saving the bound PDGFRβcells. The resulting pericytes (CD31- & PDGFRβ+) were placed in a T75 collagen coated flask and grown until confluence.

#### Immunocytochemistry

Cardiomyocytes on glass coverslips, were fixed in fresh 4% paraformaldehyde in phosphate buffered saline (PBS: 0.1 M sodium phosphate buffer, 0.9% NaCl, pH 7.4) and subsequently blocked with 5% goat serum in PBS containing 0.3% Triton X-100 for 90 min at room temperature, then incubated overnight at 4 °C with primary antibodies diluted in blocking buffer. The following primary antibodies and dilutions were used: rabbit anti-GPR39, 1:100 (Novus Biologicals, #NLS139), mouse anti-Troponin I, 1:100 (Thermo Fisher Scientific, #MA5-12960). Cells were washed with PBS + 0.1% Tween 20 and secondary antibody (1:200, Alexa 488-conjugated donkey anti-rabbit or Alexa 647-conjugated donkey anti-mouse (Thermo Fisher Scientific, Cat. #A21206 and #A31571)) were applied in blocking buffer for 2 h at room temperature. Cell nuclei were labeled with Hoechst 33342 (Thermo Fisher Cat. #62249). The coverslips were washed and mounted using Fluoromount-G® (SouthernBiotech, #0100-01). Images were acquired with a confocal microscope (Nikon Eclipse Tie-A1RSi).

#### Immunohistochemistry

Hearts were removed from rats that express Tomato red under PDGFRβ promotor (Cyagen Biosciences, Santa Clara, CA, USA). Tissues were fixed with fresh 4% paraformaldehyde and cut by cryostat into 20-µm sections. The sections were mounted on superfrost glass slides. After 90 min of blocking in 5% goat serum 1% Bovine Serum Albumin 0.3% Triton-100 in 0.1M PBS at room temperature, sections were double labelled with following primary antibodies (rabbit anti-GPR39, 1:100 (Novus Biologicals, #NLS139); mouse anti-Troponin I, 1:200 (Thermo Fisher Scientific, #MA5-12960) All primary antibodies were diluted in blocking buffer. and incubated overnight at 4 °C. All the secondary antibodies (1:400, Alexa 488-conjugated donkey anti-rabbit or Alexa 647-conjugated donkey anti-mouse (Thermo Fisher Scientific, Cat. #A21206 and #A31571)) were diluted in PBS and incubated at room temperature for 2 h. After sections were stained 2.5 min with 1:6000 Hoechst 33342 (Thermo Fisher Cat. #62249) and mounted using Fluoromount G® (SouthernBiotech, #0100-01). The sections were viewed on confocal microscope, and digital images were taken with Nikon Imaging System (Nikon Eclipse Tie-A1RSi).

#### Western Blot

Tissue from 3 hearts were homogenized in liquid nitrogen and lysed in RIPA buffer (Thermo Fisher Scientific, Cat. # PI89900) with protease and phosphatase inhibitor (Pierce, Thermo Fisher Scientific, Cat. # A32959). Cells (cardiomyocytes, pericytes and VSMCs) underwent three freeze/throw cycles in RIPA buffer with protease and phosphatase inhibitor. The homogenates were then centrifuged at 14000 rpm for 20 min at 4°C. The Pierce BCA Protein assay kit (Thermo Fisher Scientific, Cat. # 23225) was used to measure the concentration of protein lysate in samples. 50 µg/lane for cells and 20 ug/lane for whole heart tissue were separated on 4%–12% SDS-polyacrylamide gels and transferred to polyvinylidene difluoride membranes. Total protein staining was performed using Licor Bio Revert® 700 total Protein Stain (Licor Bio, Cat. #926-11016) following manufacture protocol. The membranes were blocked with Licor Intercept® TBS blocking buffer (Licor Bio, Cat. #927-60003) for 60 min at room temperature followed by primary antibodies blocking buffer with 0.02% Tween-20: 1:1,000 rabbit anti-GPR39 antibody (Millipore Sigma, Cat. #SAB4200185-200UL). After 1h incubation with secondary antibody (IRDye® 800CW Goat anti-Rabbit IgG Secondary Antibody, 1:5000, Licor Bio, Cat. # 926-32211) at room temperature, the membranes were imaged using Licor Odyssey Imager with Image Studio software. The intensity of GPR39 protein band was normalized to total protein stain in each sample. Fold change expression was calculated relative to whole rat heart sample.

#### qPCR

qPCR was performed to determine the relative quantity of GPR39 within the cell types and whole heart tissue. The relative amount of GPR39 was calculated by utilizing the Comparative Ct method (ΔΔCt) normalized to β-actin as an endogenous control. ΔΔCt values were then used to find the relative fold change in expression (2^(-(ΔΔCt))^) of each cell type in relation to the expression in the whole heart tissue.

Total RNA was recovered from isolated rat heart tissue, pericytes, VSMCs, and cardiomyocytes using RNeasy Plus Mini Kit (Qiagen, Cat. # 74104). The whole tissue was extracted after euthanasia, the thorax was opened, and the heart was immediately flash frozen in liquid nitrogen. RNA quality was inspected and verified on a NanoDrop prior to further use. Approximately 200 ng of RNA was reverse-transcribed to generate first-strand complementary DNA (cDNA) with a High-Capacity cDNA Reverse Transcription Kit with RNAse inhibitor (Thermo Fisher Scientific, Cat. # 4374966). cDNA was subjected to PCR amplification using gene-specific Taqman primers (Thermo Fisher Scientific). The 10 μL reaction containing Taqman Fast Advanced Master Mix (Thermo Fisher Scientific, Cat # 4444557), Taqman primers (Thermo Fisher Scientific, GPR39: Rn03037275_s1, β Actin: Rn00667869_m1), RNA, and nuclease free H2O. The experiments were performed by following Taqman’s instruction with Applied Biosystems Quantstudio 7 Flex Real-Time PCR System (RRID:SCR_022651).

### In vivo Experiments

#### Study protocols

Rats underwent 1 h of coronary occlusion to create AMI, after which 1 h of reperfusion was performed in all but group 5 rats. Groups 1 to 4 were randomized to drug versus vehicle and the in vivo read outs included systemic hemodynamics, tissue pO_2_, regional wall thickening (WT), INV and NRV. Figure 1 details the in-vivo study protocols indicating time points where measurements were made. Before any intervention or measurement, oxyphor was injected into the myocardium in Groups 1 to 4 animals. Group 1 animals were administered the drug (0.5 mg/kg) or vehicle 15 min prior to coronary occlusion. INV (Group 1A) and NRV (Group 1B) were measured 1 h after reperfusion. Group 2 animals received drug (0.5 mg/kg) or vehicle 15 min after coronary occlusion and INV measured 1 h after reperfusion. In Group 3 animals, VC108 (0.5 mg/kg) or vehicle, was administered 5 min prior to reperfusion to simulate the clinical scenario of attempted reperfusion, and INV (Group 3A) and NRV (Group 3B) measured 1 h after reperfusion. In a subset of animals (Group 3 C), histopathology was performed for the presence of necrosis, apoptosis, and ferroptosis. Additional Group 3B animals were used to test 3 other doses of VC108 (0.1, 0.3, and 1.0 mg/kg) for NRV reduction. In Group 4 animals, VC108 (0.5 mg/kg) or vehicle, was administered 30 min after reperfusion and INV (Group 4A) and NRV (Group 4B) measured 1 h after reperfusion. Group 5 animals received 3 different doses of VC108 15 min after coronary occlusion and animal euthanized 45 min later, at which time drug levels were measured in plasma as well as in the ischemic and normal myocardium. Finally, in Group 6 animals, 3 different doses of VC108 were administered 5 min prior to reperfusion and plasma drug levels were measured just before and several times after reperfusion, and tissue drug levels in the risk area and normal myocardium were measured 60 min after reperfusion.

**Figure 1:**
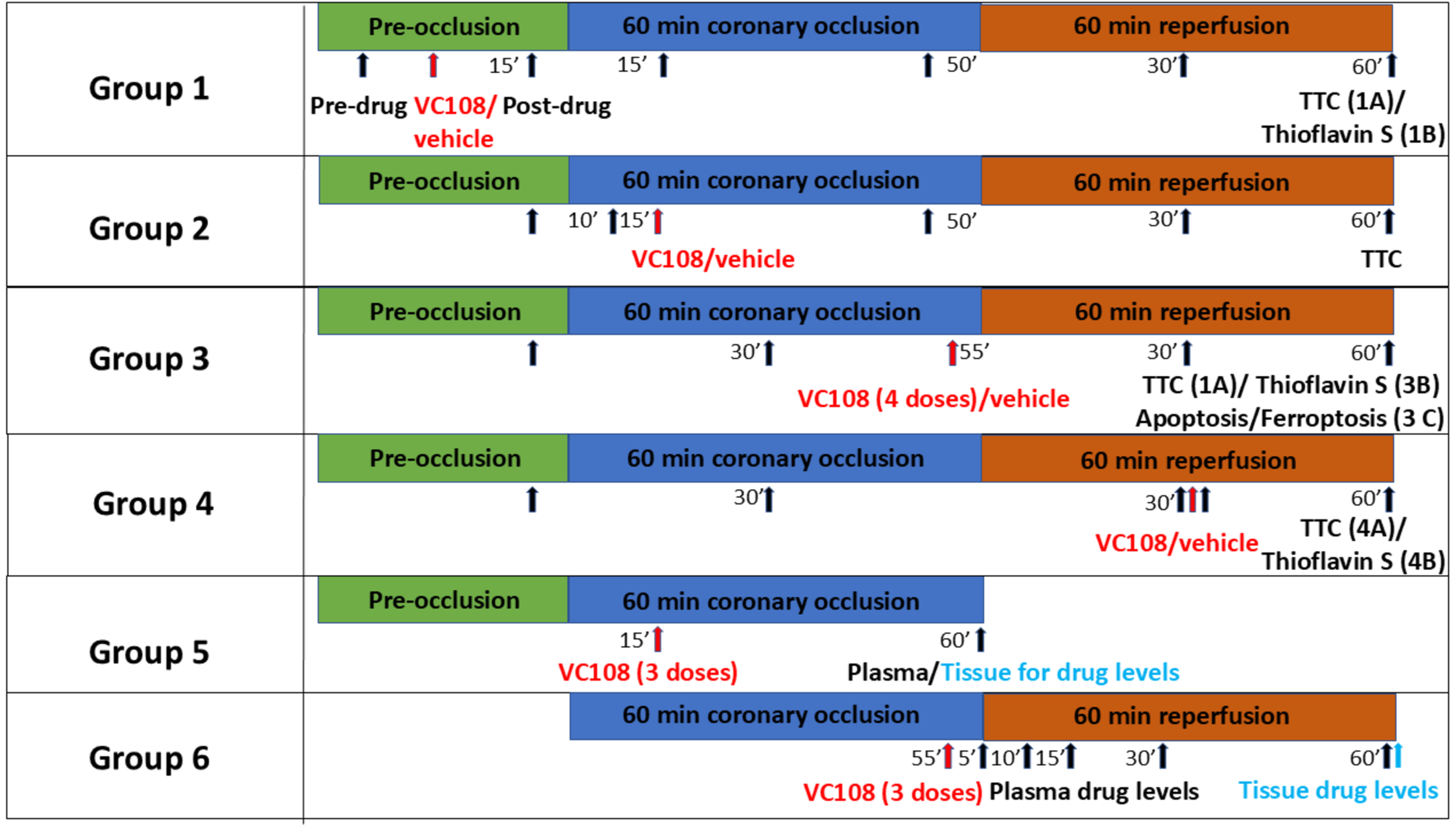
In vivo Study protocols. Groups 1 to 4 animals underwent a one-time myocardial oxyphor injection at 3-5 sites in the myocardium just after heart exposure and 30 min prior to any intervention or measurement. All animals underwent 1 hour of coronary occlusion, and all except Group 5 animals, then underwent 1 hour of reperfusion. Groups 1A and 1B underwent a similar protocol except that the final readouts were different: INV versus NRV, respectively. Both groups received 0.5 mg/kg of VC108/vehicle 15 min prior to coronary occlusion. Group 2 received 0.5 mg/kg of VC108 or vehicle 15 min into coronary occlusion and INV measured after reperfusion. Group 3 animals received VC108/vehicle 5 min prior to reperfusion, with INV measured in Group 3A and NRV measured in Group 3B where multiple doses of VC108 were given. Group 3C animals underwent histopathology for necrosis, apoptosis, and ferroptosis. Groups 4A and 4B received 0.5 mg/kg of VC108 30 min into reperfusion and INV and NRV measured, respectively. Group 5 received different doses of VC108 15 min post coronary occlusion for measurement of blood and tissue drug levels 45 min later. Group 6 received different doses of VC108 5 min before reperfusion with blood levels measured periodically up to 1 h after reperfusion at which time tissue drug levels were also measured. The black arrows indicate the timings of measurements. Groups 1 to 4 measurements included heart rate, mean aortic pressure, WT (except Group 4), and tissue pO_2_, and Groups 5 and 6 measurements included blood and tissue drug levels. Red arrows depict timing of vehicle or drug administration.

Additionally, pharmacokinetic studies of VC108 in Sprague Dawley rats was performed following a single intravenous injection in 3 males (1 mg/kg) and plasma concentrations measured over time). Then VC108 was given to 60 rats (10/sex/group) at doses of 4, 15 and 40 mg/kg/day once daily for 14 days by intravenous (bolus) injection. Dose volume was 5 mL/kg/day.

#### Acute Myocardial Infarction Creation

Rats were anesthetized with isoflurane (3% induction and 1.5% maintenance), intubated, connected to a ventilator (Harvard apparatus; Holliston, MA, USA), and placed in supine position on a warming pad to maintain a body temperature at 37°C. ECG electrodes were placed on both the arms and right leg of the animals and connected to an echocardiography system for continuous ECG monitoring. ECG was recorded at each stage during echocardiographic data acquisition. A PE20 catheter was placed in the femoral vein for administration of drug/vehicle and fluids as needed. A PE10 catheter was inserted in the left femoral artery and connected to AxonCNS Digidata 1440A interfaced with AxoScope 10.7, (Molecular Devices, San Jose, CA, USA) for continuous recording of mean aortic pressure.

The heart was exposed in the 4^th^ intercostal space and the ribs were retracted. An 8-0 ligature was placed around the left main coronary artery (LC), which was occluded by pulling the ends of the ligature through a PE10 tubing and clamping the tubing over the artery.^2^ Myocardial ischemia was confirmed by abrupt ECG changes and myocardial pallor. After 60 min of LC occlusion the PE tubing was removed and the heart reperfused for 1 h in Groups 1-4, and Group 6 animals.

Ischemic risk volume (IRV) was defined by re-occluding the LC after 1 h of reperfusion and injecting 0.2 mL of 1% Evans blue dye (Sigma Aldrich, St. Louis, MO, USA) into the aorta.^2^ NRV was defined by injecting Thioflavin-S (Sigma Aldrich) into the left atrium at the end of the reperfusion period and prior to LC re-occlusion and Evans blue dye injection.^2^ Animals in group 3C were injected with 0.1 mL of 2.0 mg/mL Propidium iodide (PI, TargetMol, Wellesley Hills, MA, USA) into the left atrium at the end of reperfusion period, followed by LC re-occlusion and Lycopersicon esculentum (Tomato) Lectin (LEL, TL), DyLight 649 (0.5 ml of 0.2 mg/mL, Thermo Fisher Scientific, Cat # L32472) injection into the aorta to define risk area. The animals were euthanized by cervical dislocation. Group 5 animals were euthanized 45 min after drug administration and heart tissue and plasma harvested for analysis. In Group 6 animals, repeated venous blood sampling was performed just prior to and several times during the 1 h of reperfusion, and then heart removed after euthanasia for drug level estimation.

#### Drug/Vehicle Administration

In groups 1 to 4 animals, either VC108 or vehicle was given as a single intravenous bolus, the timing and dose of which differed as indicated in Figure 1 and Table 1. It was dissolved in 10% dimethyl sulfoxide in saline. Total injected volume was 200 µL. The same volume of 10% dimethyl sulfoxide was injected for vehicle. Groups 3B and Group 5 animals received the same volume of drug but at different doses (Table 1). Group 6 animals received 0.33 mL/kg for 0.1 mg/kg dose, 0.5 mL/kg for 0.3 mg/kg dose, and 0.75 mL/kg for 1.0 mg/kg dose. VC108 is a selective GPR39 antagonist.^12^ Its in vitro absorption, distribution and metabolism properties are described in Supplemental Table 1.

**Table 1.** Completed Animals Used for the In-vivo Study (n=280)

| **Group** | **Final Readout** | **Timing of Drug/**<br>**Vehicle IV** | **Intervention** | **Animal**<br>**Number** | **Animal**<br>**sex** |
| --- | --- | --- | --- | --- | --- |
| **1A** | Infarct Volume | 15 min Before Coronary Occlusion. | Vehicle (control)<br>VC108 0.5 mg/kg | 11<br>16 | 6 M, 5 F<br>9 M, 7 F |
| **1B** | No Reflow Volume | 15 min Before Coronary Occlusion. | Vehicle (control)<br>VC108 0.5 mg/kg | 11<br>11 | 6 M, 5 F<br>6 M, 5 F |
| **2** | Infarct Volume | 15 min After Coronary Occlusion. | Vehicle (control)<br>VC108 0.5 mg/kg | 14<br>12 | 7 M, 7 F<br>6 M, 6 F |
| **3A** | Infarct Volume | 5 min Before Reperfusion. | Vehicle (control)<br>VC108 0.5 mg/kg | 15<br>13 | 9 M, 6 F<br>6 M, 7 F |
| **3B** | No Reflow Volume | 5 min Before Reperfusion. | Vehicle (control)<br>VC 108 0.1 mg/kg<br>VC108 0.3 mg/kg<br>VC108 0.5 mg/kg<br>VC108 1.0 mg/kg | 15<br>15<br>11<br>13<br>12 | 8 M, 7 F<br>7 M, 8 F<br>5 M, 6 F<br>6 M, 7 F<br>6 M, 6 F |
| **3C** | Apoptosis/<br>Ferroptosis | 5 min Before Reperfusion | Vehicle (control)<br>VC108 0.5 mg/kg | 6<br>6 | 3 M, 3 F<br>3 M, 3 F |
| **4A** | Infarct Volume | 30 min after Reperfusion. | Vehicle (control)<br>VC108 0.5 mg/kg | 19<br>19 | 11 M, 8 F<br>11 M, 8 F |
| **4B** | No Reflow Volume | 30 min after Reperfusion. | Vehicle (control)<br>VC108 0.5 mg/kg | 15<br>19 | 7 M, 8 F<br>9 M, 10 F |
| **5** | Plasma/<br>Tissue drug level | 45 min After Coronary Occlusion. | VC108 0.5 mg/kg<br>VC108 2.5 mg/kg<br>VC108 5.0 mg/kg | 6<br>6<br>6 | 3 M, 3 F<br>3 M, 3 F<br>3 M, 3 F |
| **6** | Plasma/<br>Tissue drug levels | Plasma: before and 5,10,15,30, and 60 min after reperfusion.<br>Tissue: 60 min after reperfusion. | VC108 0.1 mg (0.33 mL)/kg<br>VC108 0.3 mg (0.5 mL)/kg<br>VC108 1.0 mg (0.75 mL)/kg | 3<br>3<br>3 | 3 M<br>3 M<br>3 M |

M=male; F=female; IV=intravenous (bolus)

#### Tissue Oxygen Tension

Tissue pO_2_ was measured as an indicator of tissue perfusion within the IRV.^16^ An OxyLED TD phosphorometer (Oxygen Enterprises, Havertown, PA, USA) connected to optical fibers for excitation and collection of emitted phosphorescence was used. A phosphorescence probe (Oxyphor G4, 0.3 ml of a 200 M solution; Oxygen Enterprises) derived from the Pd-meso-tetra-(3,5-dicarboxyphenyl) tetrabenzoporphyrin that does not permeate biological membranes and is highly soluble in aqueous environments was used.^17^ Three to five intramyocardial injections were made into the anterolateral surface of the heart (exposed area) 30 min prior to the first measurement. The light guide’s tips were placed 2-3 mm from the heart surface for tissue O_2_ tension measurement. The Stern-Volmer relationship was used to convert the measured phosphorescence lifetimes to pO_2_.^18^

#### Wall Thickening Measurements

Regional WT was measured using 2-dimensional echocardiography with a Sonos 5500 system (Philips, Cambridge, MA, USA) interfaced to a S12 transducer that transmits ultrasound between 5 and 12 MHz frequencies. Imaging was performed in the parasternal short-axis view incorporating the papillary muscle with the transducer affixed to the procedure table. Ultrasound gel was placed between the heart and transducer. We identified the post-mortem tissue slice closest to the echocardiographic slice using landmarks such as the papillary muscle and insertion points of the right ventricular free wall to the left ventricle. Echocardiographic analysis was only performed if a risk area was present in the slice that approximated the echocardiographic image.

WT was measured by a single observer blinded to all other data.^19^ The epicardial junction of the right ventricular posterior wall and the LV free wall was identified in each frame for image registration. A dozen targets were then outlined on the endocardium and epicardium, respectively, in each frame from end-diastole to end-systole, which were then automatically connected using cubic spline interpolation. One hundred equidistant points were automatically defined on the epicardial outline to which a tangent was drawn and a line perpendicular to the tangent intersected the epicardium and endocardium as the shortest distance between them. The resulting chord lengths were measured in all frames from end-diastole to end-systole with the first chord being the reference point. From these WT was calculated within the pre-defined regions.

#### Gross Tissue Staining

All measurements were made by a single observer blinded to all other data.^2^ After euthanasia, the heart was removed and cut into 5-6 cross sections. Left ventricular (LV) volume was first calculated by planimetering the epicardial and endocardial outlines of the LV in each slice and excluding the right ventricle, then summing up the LV areas from all slices. For IRV assessment regions not stained blue were planimetered in each slice and their values summed. IRV was expressed as percent of LV myocardial volume.

For NRV, heart slices were photographed under Ultraviolet light (420 nM wavelength) where Thioflavin-S fluoresces blue.^2^ NRV was calculated using two methods. First, LV regions devoid of Thioflavin-S were planimetered and summed, then expressed as percent of the LV IRV. Second, all images were converted to 8-bit gray scale images and the severity of no reflow defined as ratio of mean pixel intensity in the no reflow zone versus pixel intensity in normal myocardium. This fraction was then multiplied by the planimetered volume. The second measurement was performed because of variation in pixel intensity in the no reflow zone.

INV assessment was performed by staining heart slices with 1% 2,3,5 triphenyl tetrazolium chloride (TTC, Sigma Aldrich) in phosphate buffered saline for 20 min at 37°C followed by incubation in 10 % neutral buffered formalin overnight.^2^ The sections were imaged using a digital camera. The infarcted region, unstained by TTC was planimetered and normalized to the IRV.

#### Measurement of Necrosis, Apoptosis, and Ferroptosis

Hearts were fixed in fresh 4% PBS and paraffin sections of 6 um thickness were placed on slides for assessing necrosis, apoptosis, and ferroptosis.^20^ For assessing necrosis, propidium iodide was injected into the left atrium at the end of reperfusion period. Paraffin sections were deparaffinized and rehydrated followed by washing once with PBS-T and stain with Hoechst (cat #33342) for DAPI (1:6000 of 10mg/ml stock for 2.5 min) and washed in PBS before mounting using Fluoromount-G (Southern Biotech).

For assessing apoptosis, deparaffinized and rehydrated sections were placed in 10 mM sodium citrate buffer (pH 6.0) at a sub-boiling temperature for 30 min. After cooling the slides, a 30 min antigen retrieval was initiated. The slides were washed in PBS and then immersed in PBS containing 5% normal goat serum, 1% bovine serum albumin, and 0.3% Triton X-100 for 90 min at room temperature. The sections were then incubated overnight at 4 °C with primary antibody 1:100 Annexin V (Cell signaling, Cat: #54043T). Secondary antibody, 488 goat anti rabbit (1:400, Thermo Fisher, Cat: #A11008) was incubated for 2 h followed by DAPI stain.

For assessing ferroptosis, the heart slides were deparaffinized and rehydrated, followed by incubation for 3 min using an Iron stain Kit (Abcam, Cat #AB150674) containing a mixture of potassium ferrocyanide and hydrochloric acid. The slides were then rinsed in distilled water for 4 min before counterstaining with nuclear fast red for 5 min. After another wash with distilled water, slides were dehydrated for 5 min in alcohol and CitriSolv (Decon Labs. Inc., Cat #1601) and mounted using Cytoseal -60 (Thermo Fisher Scientific. Cat: #8310-4).

All slides were imaged using Zeiss Axio Scan 7 confocal slide scanning microscope. One experienced, blinded observer then read the slides for presence of iron (bright blue), PI (red) and Annexin V (green) on a scale of 0 to +4.

#### Plasma and Tissue Drug Measurement

In Group 5 rats, after euthanasia, 1 mL of blood was collected by cardiac puncture. In Group 6 animals, venous blood was withdrawn just before and at different times after reperfusion. In both groups of animals, the heart was removed at the end of the experiment and atria and aorta discarded. The left ventricle was divided into the pale occluded part and the normal myocardium. Samples were stored in liquid nitrogen and transported to Aptuit (Verona, Italy) for analysis.

A 12-point calibration curve and quality control samples were prepared in plasma diluted 1:4.8 in 0.1N Hepes buffer in micronic tubes. Twenty µL of plasma was transferred into micronic tubes containing 80µL 0.1N Hepes buffer. Heart samples were diluted 1:5 in 0.1N Hepes buffer and homogenated by Precellys tissue homogenizer (Bertin Technologies, Montigny-le-Bretonneux, France). The homogenate (100µL) was then transferred into micronic tubes. Calibration samples (CS), quality control samples (QC) and analysis samples were extracted by adding 400µL CH3CN containing IS (Rolipram) 20 ng/mL.

After vortex and centrifugation (3000 rpm/10 min), samples supernatants were transferred to a 96 well plate and diluted (450µL CH3CN/aq=37/63 + 50µL supernatant). Samples were injected using an Agilent 1290 Infinity II UPLC system (Santa Clara, CA, USA) and separated on a Acquity UPLC BEH C18 (30×2.1mm, 1.7µm, Milford, Massachusetts) connected directly to the TurboIonSpray source of an AB Sciex 6500QTrap mass spectrometer (Framingham, MA, USA). Mobile Phase A was 0.1% (v/v) formic acid in water and Mobile Phase B was 0.1% (v/v) formic acid in acetonitrile. The flow rate was 1 mL/min and a generic gradient of 5% B to 95% B in 1.3 min was applied.

MS Conditions
| Analyte | Precursor ion (m/z) | Product Ion (m/z) | Polarity |
| --- | --- | --- | --- |
| APV239108A | 415.2 | 294.3 | Positive |
| Rolipram (IS) | 276.1 | 208.2 | Positive |

### MS Conditions

To determine free plasma fraction of drug, 0.5 µM of plasma samples (in triplicate) were loaded onto a 96-well equilibrium dialysis apparatus (HTDialysis, Gales Ferry, CT, USA) and dialyzed against an equal volume of buffer at 37°C. Positive controls were included. After 5 h aliquots of plasma and buffer were transferred to a 96-deep well plate and the composition in each tube was balanced with control fluid making the volume of each well the same. Sample extraction was performed by adding acetonitrile containing an internal standard. Samples were analyzed in a LC-MS/MS to monitor compound to internal standard peak area ratios as representing compound concentrations.

#### Statistical Methods

Power calculation was performed for estimation of sample size, where drug was estimated to result in 30% reduction in NRV or INV as percent of IRV. Unless otherwise noted, data are expressed as mean ± 1 standard error of the mean (SEM). Male and female results were first considered separately. If no differences were found, then they were presented together. Comparisons between two data sets were performed using unpaired nonparametric Mann-Whitney test followed by Kolmogorov-Smirnov post hoc test. More than 2 data sets from the same animals were compared using 2-way analysis of variance (ANOVA) with repeated measures. When ANOVA showed significant differences, selected groups were compared using paired *t t*est and p value adjusted for multiple comparisons. Results were considered significant at *P*<0.05.

## Results

### In vitro Experiments

Figure 2A illustrates ICC of cardiomyocytes stained with DAPI (blue), Troponin I (green), and GPR39 (magenta). The merged image shows colocalization of GPR39 with Troponin I (scale bar = 20 µm). Figure 2B illustrates the results of qPCR of whole rat cardiac tissue (control, n=6) as well as isolated primary cardiomyocytes (n=5), pericytes (n=5), and VSMCs (n=5) (all normalized to β-actin). Cardiomyocytes express GPR39 at the same level as VSMCs.

**Figure 2:**
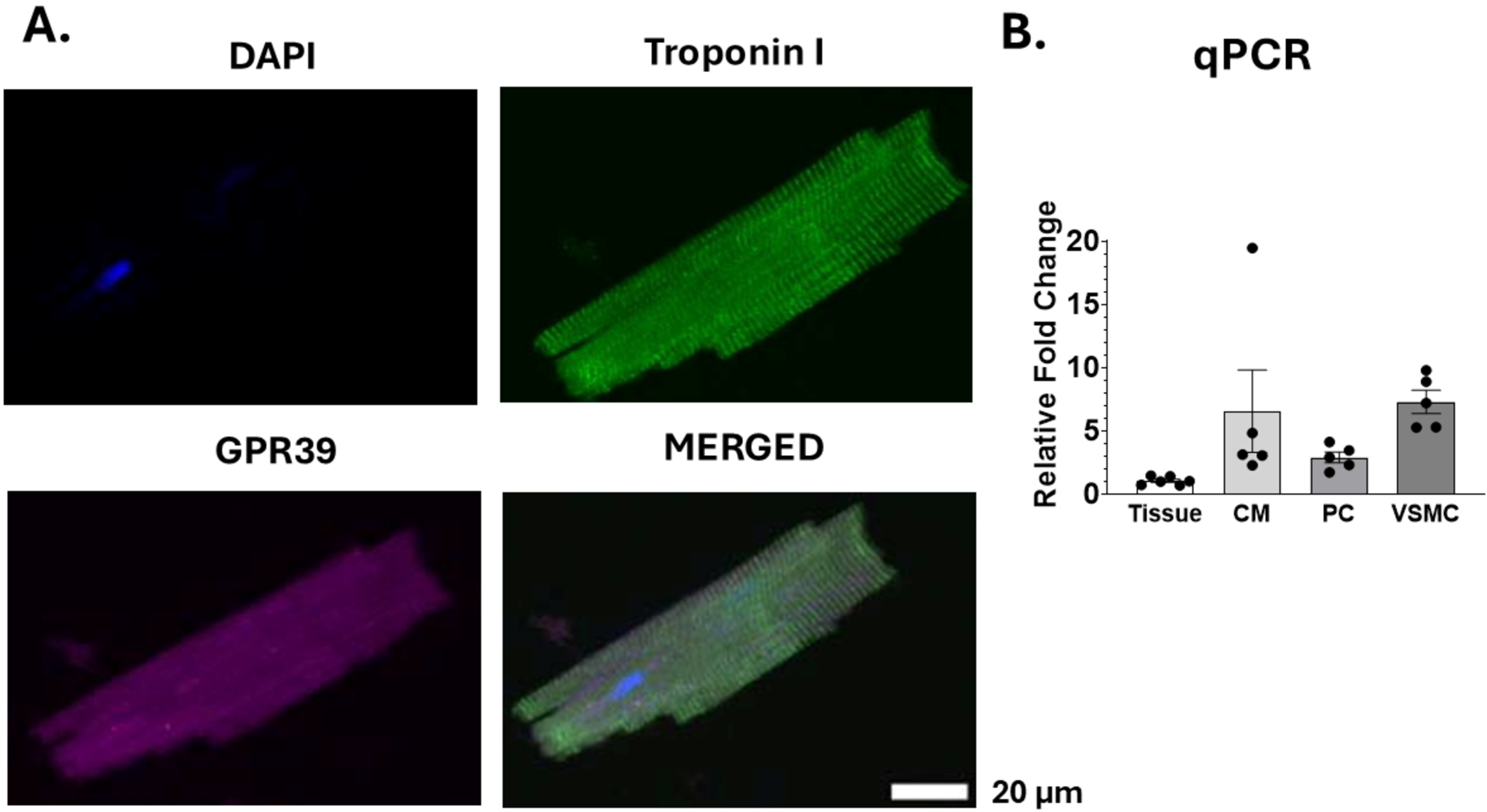
Results of ICC and qPCR. (A) ICC of an isolated rat cardiomyocytes showing DAPI (blue), Troponin (green), and GPR39 (magenta) along with merged image. (B) qPCR results from rat cardiac tissue (control, n=6), cardiomyocytes (n=5), pericytes (n=5), and VSMCs (n=5) (normalized to β-actin, scale bar = 20 µm), showing almost equivalent GPR39 gene expression in cardiomyocytes in comparison to VSMCs, which is higher than that in pericytes.

Figure 3A depicts IHC of rat myocardial tissue demonstrating staining of nuclei (DAPI, blue), pericytes and VSMCs (PDGFRβ, red), cardiomyocytes (troponin I, magenta) and GPR39 (green). The merged image shows co-localization of all staining (scale bar = 20 µm). Figure 3B illustrates WB from a representative sample along with aggregate data (n=3) shows presence of GPR39 in all cell types. GPR39 protein expression in cardiomyocytes is less than the mRNA expression.

**Figure 3:**
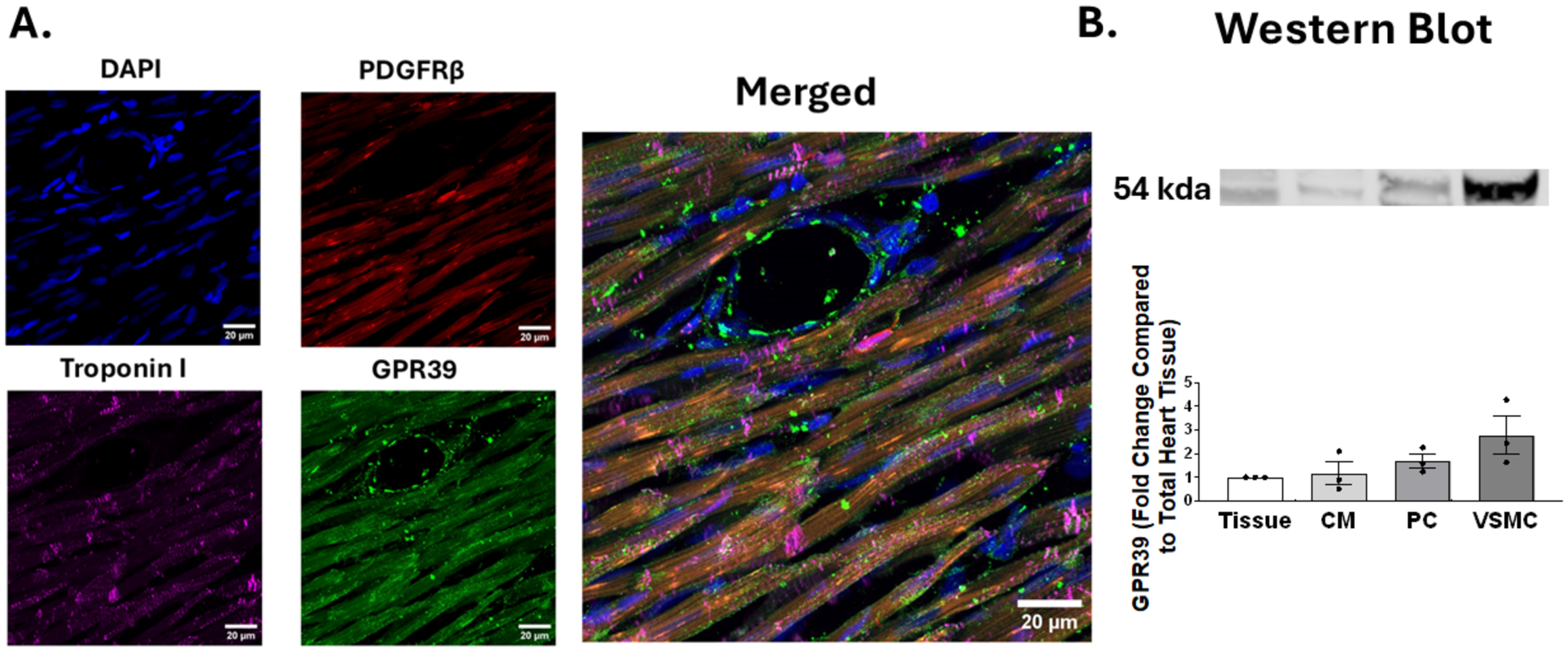
Results of IHC and Western Blot. IHC of rat myocardial tissue demonstrating nuclei (DAPI, blue), pericytes and VSMCs (PDGFRβ, red), cardiomyocytes (troponin I, magenta) and GPR39 (green). The merged image shows co-localization of all staining (scale bar = 20 µm). Figure 3B illustrates a WB sample showing presence of GPR39 protein in all cell types with aggregate results from 3 samples.

### In vivo studies

Of a total of 357 animals undergoing coronary occlusion, 280 survived the study duration (22% mortality). Majority of the 40 males and 37 females died from ventricular fibrillation or ventricular tachycardia within 15 min of coronary occlusion; a minority had bradycardia. The number and sex of the surviving animals from each group is shown in Table 1. There were no differences in the results between males and females in any of the readouts from Groups 1 to 5. Group 6 comprised only males. Hence, the aggregate results represent the entire cohort although in the bargraphs males and females are shown in different colors (black males, red females).

### Groups 1 Animals

These animals were administered either VC108 (0.5 mg/kg) or vehicle before coronary occlusion. Hemodynamics, tissue pO_2_, and WT, were measured before and after drug/vehicle administration, as well as 15 and 30 min into coronary occlusion and 30 min into reperfusion. The results from these readouts are listed in Table 2. There was no difference in heart rate between any of the stages in both vehicle and VC108 treated animals. In contrast, mean aortic pressure declined significantly during coronary occlusion in vehicle treated but not VC108 treated animals. WT declined significantly and to the same extent in vehicle and VC108 treated animals during coronary occlusion and rebounded to near normal levels in both vehicle and VC108 treated animals during reperfusion. Pre-treatment tissue pO_2_ tended to be higher in the VC108 compared to the vehicle treated group, which reversed after drug treatment. pO_2_ declined equally in both vehicle and VC108 treated animals during coronary occlusion and recovered equally in the two groups during reperfusion.

**Table 2.** Results (mean ± 1SEM) From Group 1 Animals Receiving 0.5mg/kg VC108 (n=27) or Vehicle (n=22) Prior to Coronary Occlusion.

| Variable | Pre-occlusion |  | Occlusion | Reperfusion | *P* value |
| --- | --- | --- | --- | --- | --- |
|  | Pre-treatment | Post-treatment |  |  |  |
| **Heart Rate (bpm)** |  |  |  |  |  |
| **Vehicle** | 355 ± 10* | 346 ± 10 | 344 ± 9* | 340 ± 9 | 0.4030 |
| **VC108** | 331 ± 9* | 328 ± 10 | 330 ± 9* | 331 ± 9 | 0.9278 |
| ***P* value** | 0.0671 | 0.2697 | 0.2316 | 0.5036 |  |
| **Mean Aortic Pressure (mm Hg)** |  |  |  |  |  |
| **Vehicle** | 92 ± 4* | 93 ± 4 | 80 ± 3* | 75 ± 2 | 0.0149 |
| **VC108** | 89 ± 3* | 89 ± 3 | 81 ± 3* | 69 ± 3 | 0.0875 |
| ***P* value** | 0.5737 | 0.4778 | 0.7317 | 0.0744 |  |
| **Wall Thickening (%)** |  |  |  |  |  |
| **Vehicle** | 37.6 ± 0.5 | 38.4 ± 1.8 | 15.4 ± 2.4 | 34.5 ± 0.9 | <0.001 |
| **VC108** | 37.6 ± 0.3 | 38.0 ± 1.6 | 15.7 ± 2.8 | 34.7 ± 1.4 | <0.001 |
| ***P* value** | 0.9927 | 0.8913 | 0.9543 | 0.9344 |  |
| **pO<sub>2</sub> (mm Hg)** |  |  |  |  |  |
| **Vehicle** | 32.9 ± 1.9* | 38.0 ± 1.2 | 9.1 ± 1.7* | 30.3 ± 2.0 | <0.0001 |
| **VC108** | 37.3 ± 2.0* | 34.2 ± 1.3 | 9.4 ± 1.6* | 33.1 ± 1.8 | <0.0001 |
| ***P* value** | 0.0629 | 0.0651 | 0.9121 | 0.2886 |  |

\*Comparison using paired *t*-test between groups in the row with *P* value shown in last column. Comparison only performed when repeat measured ANOVA showed $p < 0.001$ . *P* value in columns for comparison between vehicle and VC108 using unpaired *t*-test.

Figure 4 illustrates examples of IRV and INV from Group 1A animals receiving vehicle (panel A) versus VC108 (0.5 mg/kg, panel B) prior to 1 h of coronary occlusion. The INV 60 min after reperfusion is significantly smaller in the VC108 versus vehicle treated animals. Panel C depicts the aggregate results from Group 1A animals, where the INV/IRV ratio is significantly smaller with VC108 treated compared to vehicle treated animals. Panel D shows that tissue pO_2_ during coronary occlusion does not positively or negatively influence INV, being similar in the VC108 and vehicle treated animals. The relatively high pO_2_ in vehicle treated animals may be due to high collateral flow in this group during coronary occlusion.

**Figure 4:**
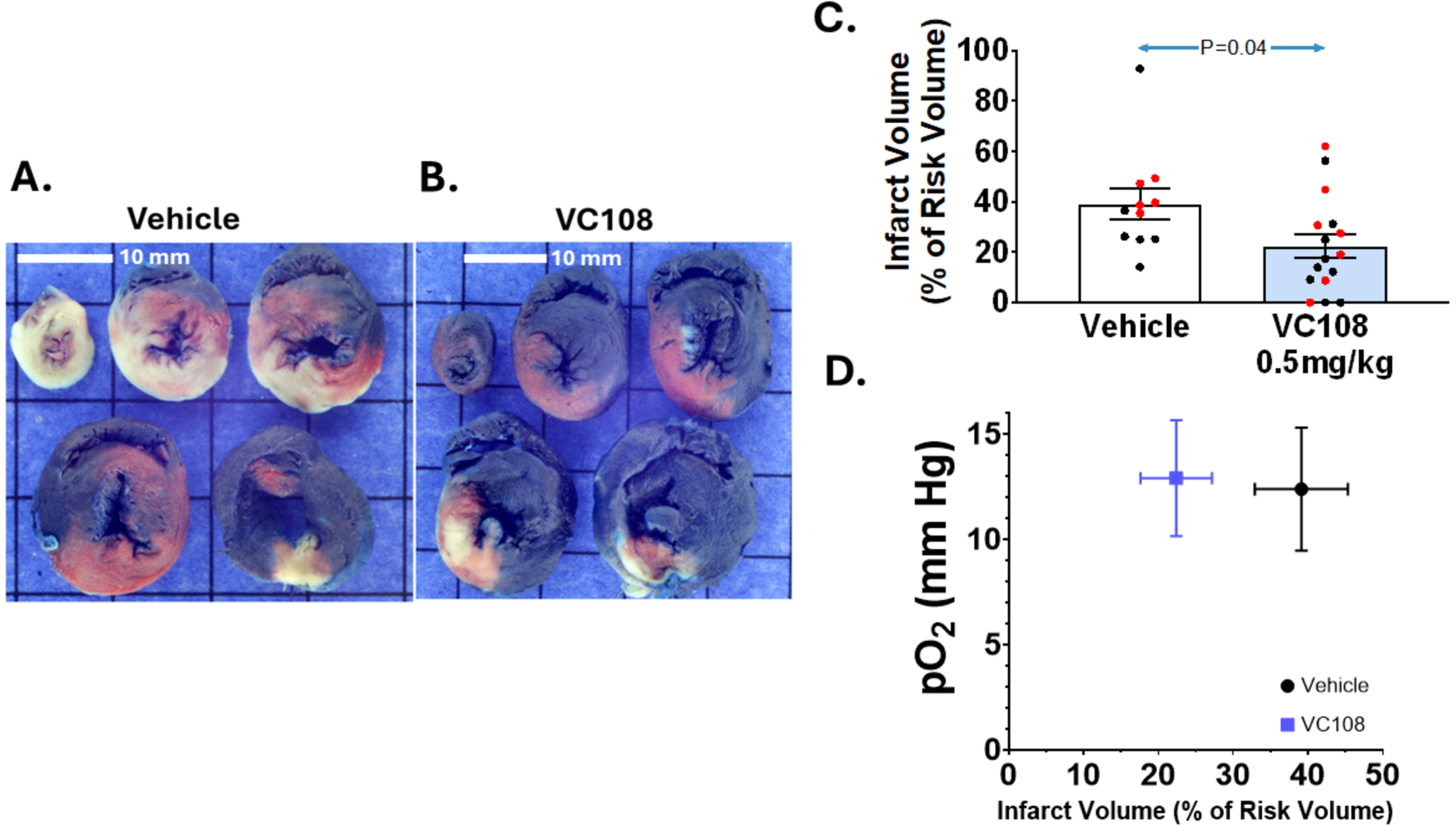
Results from Group 1A animals. Examples of INV (pale tissue) within IRV (tissue not stained blue) from a Group 1 A animal receiving either (A) vehicle (n=11) or (B) 0.5 mg/kg of VC108 (n=16) prior to coronary occlusion (scale = 10 mm). Aggregate data (C) show significantly smaller INV/IRV in VC108 treated versus vehicle treated animals. Males are depicted in black and females in red. There is no influence of tissue pO_2_ on the INV/IRV ratio (D). Differences between mean were analyzed using unpaired t-test.

Figure 5 illustrates examples of IRV and NRV from a Group 1B vehicle treated (panel A) versus a VC108 treated (0.5 mg/kg, panel B) animal where both were administered prior to coronary occlusion. The NRV is significantly smaller in the VC108 versus vehicle treated animal 60 min after reperfusion. Panel C depicts the aggregate results from Group 1B animals where the NRV/IRV ratio is significantly smaller with VC108 treated compared to vehicle treated animals. Unlike Group 1A animals, however, the results are positively influenced by tissue pO_2_ during coronary occlusion, being significantly higher in VC108 versus vehicle treated animals.

**Figure 5:**
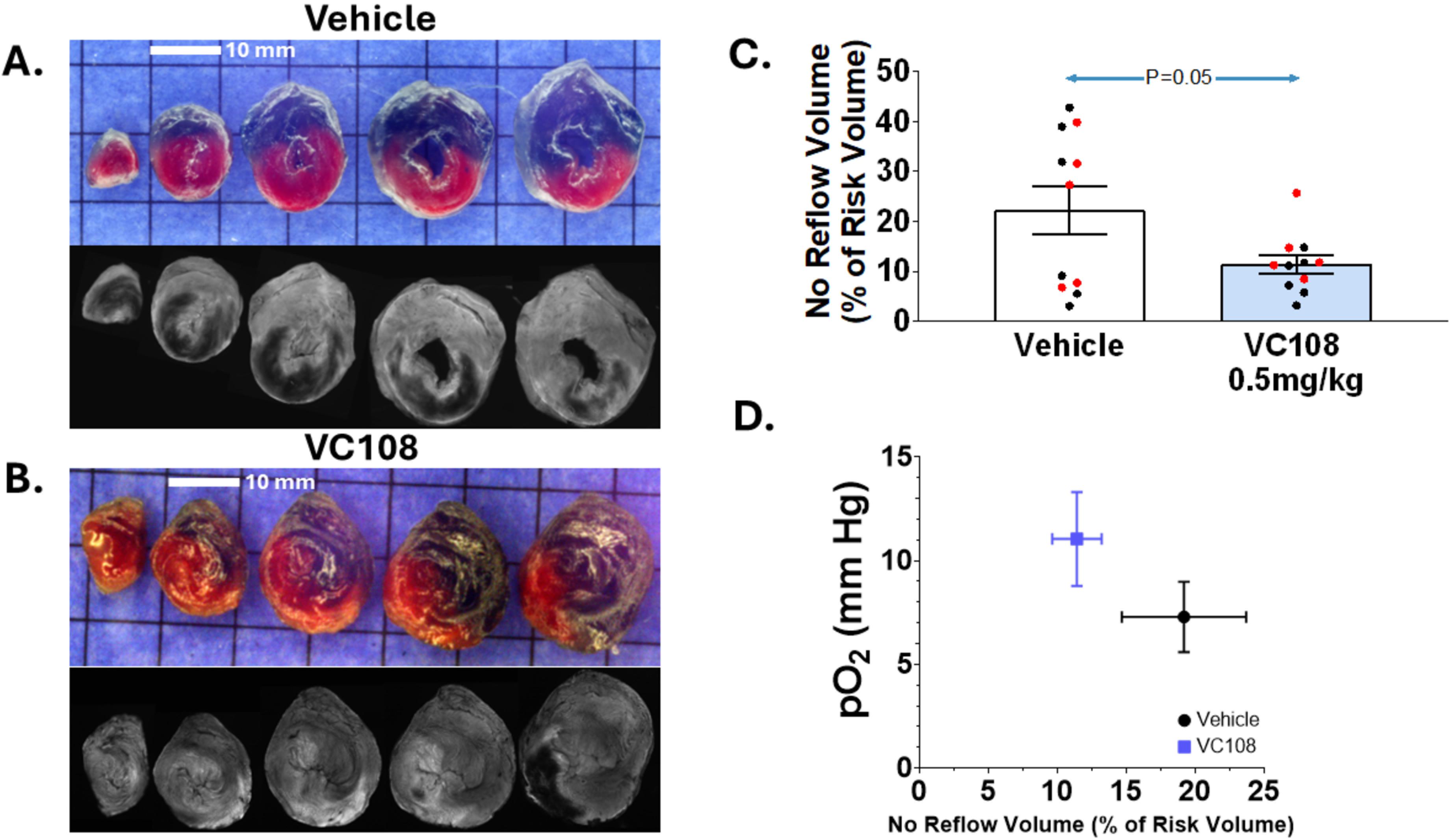
Results from Group 1B animals. Examples of no NRV (black regions) within IRV (regions not stained blue) from Group 1B animals receiving either (A) vehicle or (B) 0.5 mg/kg of VC108 prior to coronary occlusion (scale = 10 mm). Aggregate data (C) show significantly smaller NRV/IRV in VC108 treated (n=11) versus vehicle treated (n=11) animals. Males are depicted in black and females in red. Unlike in Figure 2, there is a positive influence of tissue pO_2_ on the NRV/IRV ratio. Differences between mean were analyzed using unpaired t-test.

### Group 2 animals

Table 3 depicts results from the Group 2 animals who received 0.5 mg/kg or VC108 15 min into 1 h of coronary occlusion and in which INV was measured 60 min after reperfusion. Heart rate and mean aortic pressure did not change between the stages and were not different at any stage between vehicle and VC108. WT declined precipitously and equally between vehicle and VC108 during coronary occlusion and recovered to nearly normal levels during reperfusion. Similar effects were seen with pO_2_, which was not different between pre-occlusion and reperfusion for both vehicle and VC108. However, it increased significantly during coronary occlusion after VC108 but not vehicle. pO_2_ during coronary occlusion was also significantly higher with VC108 than vehicle.

**Table 3.** Results (mean ± 1SEM) from Group 2 Animals Receiving 0.5mg/kg/ VC108 (n=12) or vehicle (n=14) 15 min After Coronary Occlusion.

| **Variable** | **Pre-**<br>**Occlusion** | **Occlusion**<br>**Pre-**<br>**treatment** | **Occlusion**<br>**Post-**<br>**treatment** | **Reperfusion** | ***P* value** |
| --- | --- | --- | --- | --- | --- |
| **Heart Rate (bpm)** |  |  |  |  |  |
| **Vehicle** | 336 ± 6 | 348 ± 12* | 346 ± 9* | 340 ± 14 | 0.9385 |
| **VC108** | 353 ± 10 | 354 ± 10* | 368 ± 16* | 358 ± 15 | 0.4656 |
| ***P* value** | 0.1621 | 0.6994 | 0.2508 | 0.4070 |  |
| **Aortic Pressure**<br>**(mm Hg)** |  |  |  |  |  |
| **Vehicle** | 78 ± 5 | 71 ± 6* | 70 ± 6* | 66. ± 4 | 0.9848 |
| **VC108** | 87 ± 3 | 70 ± 5* | 69 ± 4* | 73 ± 2 | 0.8432 |
| ***P* value** | 0.1197 | 0.9293 | 0.8079 | 0.2094 |  |
| **Wall Thickening (%)** |  |  |  |  |  |
| **Vehicle** | 35.2 ± 5.8 | 15.3 ± 7.7 | 15.1 ± 7.9 | 35.9 ± 11.1 | <0.0001 |
| **VC108** | 37.2 ± 2.1 | 16.1 ± 6.5 | 16.5 ± 6.6 | 33.1 ± 5.7 | <0.0001 |
| ***P* value** | 0.3092 | 0.7728 | 0.6545 | 0.4675 |  |
| **pO<sub>2</sub> (mm Hg)** |  |  |  |  |  |
| **Vehicle** | 30.6 ± 2.1 | 7.1 ± 1.2* | 7.4 ± 1.3* | 30.7 ± 1.5 | 0.8951 |
| **VC108** | 31.9 ± 2.5 | 5.7 ± 1.7* | 12.1 ± 1.9* | 33.1 ± 3.2 | 0.0210 |
| ***P* value** | 0.6963 | 0.5048 | 0.0473 | 0.5029 |  |

\*Comparison using paired *t*-test between pre- and post-treatment during coronary occlusion with *P* value shown in last column. *P* value in columns for comparison between vehicle and VC108 using unpaired *t*-test.

Figure 6 demonstrates IRV and INV from a Group 2 animal receiving vehicle (panel A) and VC108 (0.5 mg/kg, panel B) during coronary occlusion. The INV is markedly less in the drug treated animal. Aggregate results from Group 3 animals show a significantly smaller INV/IRV ratio in VC108 treated animals (panel C). Tissue pO_2_ during coronary occlusion positively influences the INV/IRV ratio after reperfusion, being significantly higher in VC108 treated than in the vehicle treated animals (panel D).

**Figure 6:**
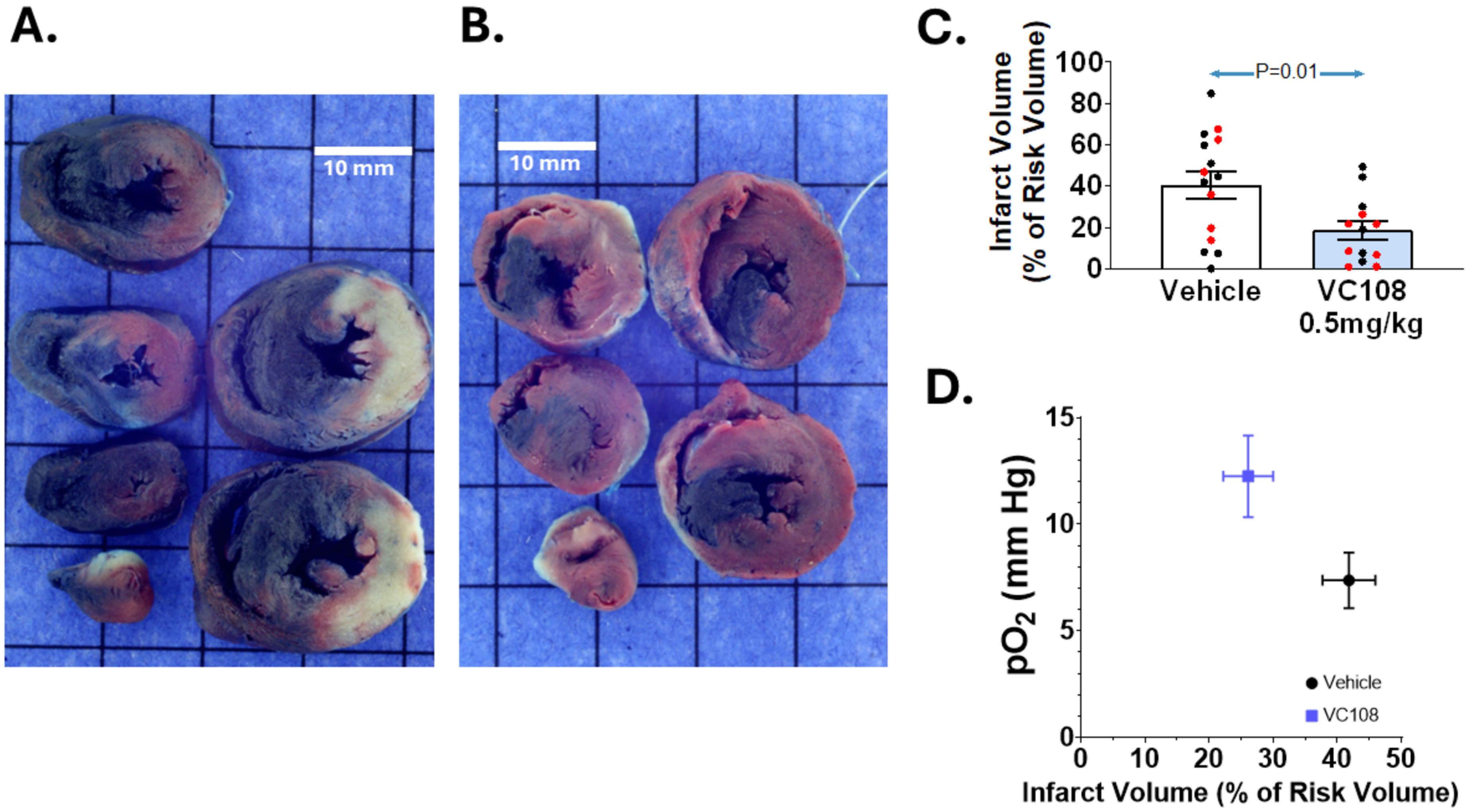
Results from Group 2 animals. Examples of INV (pale tissue) within IRV (tissue not stained blue) from Group 2 animals receiving either (A) vehicle or (B) 0.5 mg/kg of VC108 15 min into coronary occlusion (scale = 10 mm). Aggregate data (C) show significantly smaller INV/IRV in VC108 treated (n=12) versus vehicle treated (n=14) animals. Males are depicted in black and females in red. A positive influence of tissue pO_2_ on the INV/IRV ratio is seen (D). Differences between mean were analyzed using unpaired t-test.

### Groups 3 Animals

These animals received VC108 or vehicle 5 min prior to reperfusion in order to simulate the clinical scenario. They had hemodynamics, pO_2_, and WT measured at baseline, 30 min into coronary occlusion, and 30 min into reperfusion. The results from animals receiving vehicle and 0.5 mg/kg of VC108 are listed in Table 4. Heart rate did not differ between stages and between the VC108 and vehicle at each stage. The mean aortic pressure declined after coronary occlusion and remained so after reperfusion in both vehicle and VC108 treated animals with no difference between the two. Both WT and pO2 declined dramatically during coronary occlusion. WT recovered to near normal levels after reperfusion but pO_2_ did not fully recover, although it tended to be higher in the VC108 compared to vehicle treated animals.

**Table 4.**
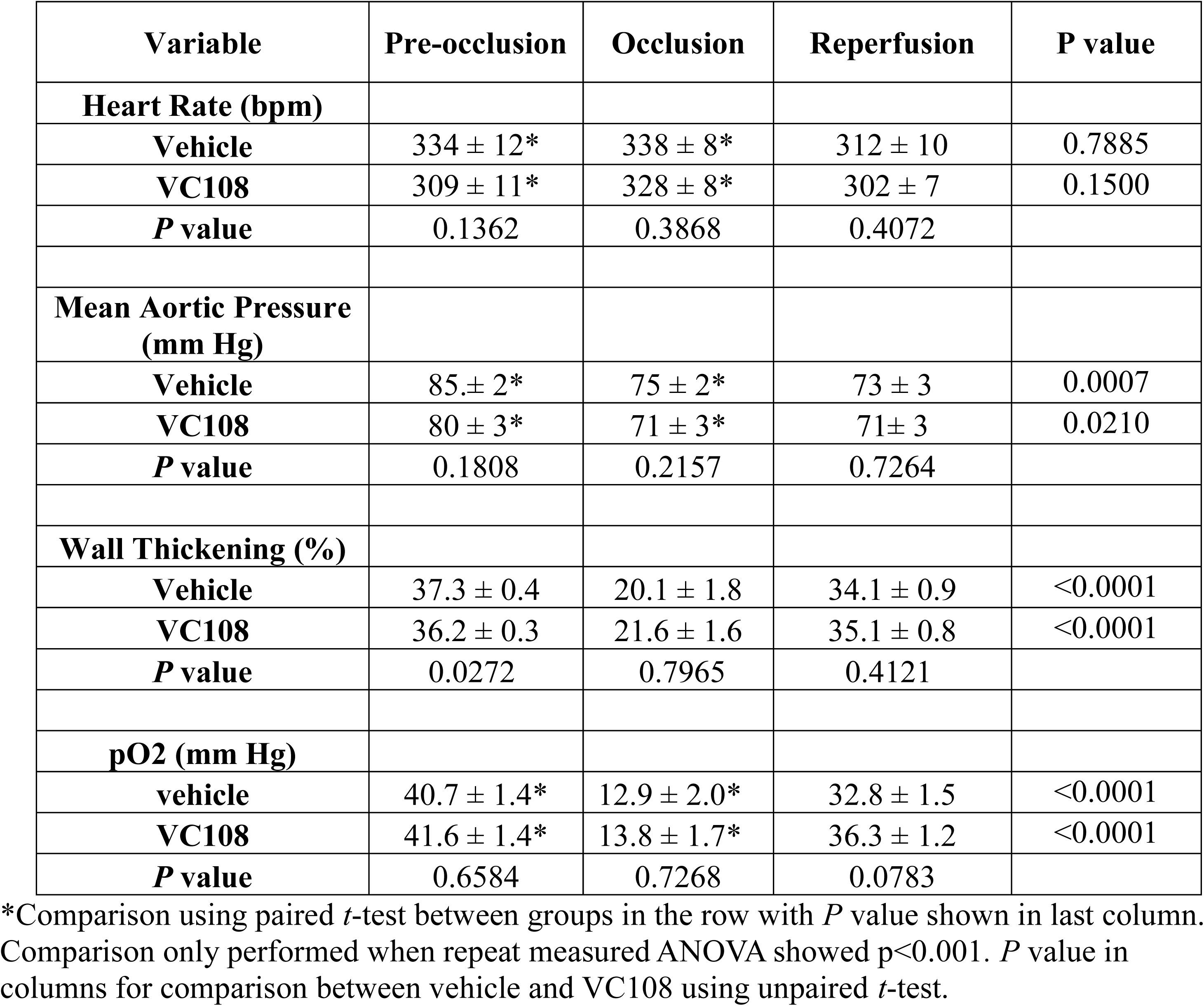
Results (mean ± 1SEM) From Group 3 Animals Receiving 0.5mg/kg VC108 (n=27) or Vehicle (n=29) 5 min Prior to Reperfusion.

| **Variable** | **Pre-occlusion** | **Occlusion** | **Reperfusion** | **P value** |
| --- | --- | --- | --- | --- |
| **Heart Rate (bpm)** |  |  |  |  |
| **Vehicle** | 334 ± 12* | 338 ± 8* | 312 ± 10 | 0.7885 |
| **VC108** | 309 ± 11* | 328 ± 8* | 302 ± 7 | 0.1500 |
| **P value** | 0.1362 | 0.3868 | 0.4072 |  |
| **Mean Aortic Pressure (mm Hg)** |  |  |  |  |
| **Vehicle** | 85. ± 2* | 75 ± 2* | 73 ± 3 | 0.0007 |
| **VC108** | 80 ± 3* | 71 ± 3* | 71 ± 3 | 0.0210 |
| **P value** | 0.1808 | 0.2157 | 0.7264 |  |
| **Wall Thickening (%)** |  |  |  |  |
| **Vehicle** | 37.3 ± 0.4 | 20.1 ± 1.8 | 34.1 ± 0.9 | <0.0001 |
| **VC108** | 36.2 ± 0.3 | 21.6 ± 1.6 | 35.1 ± 0.8 | <0.0001 |
| **P value** | 0.0272 | 0.7965 | 0.4121 |  |
| **pO2 (mm Hg)** |  |  |  |  |
| **vehicle** | 40.7 ± 1.4* | 12.9 ± 2.0* | 32.8 ± 1.5 | <0.0001 |
| **VC108** | 41.6 ± 1.4* | 13.8 ± 1.7* | 36.3 ± 1.2 | <0.0001 |
| **P value** | 0.6584 | 0.7268 | 0.0783 |  |

\*Comparison using paired *t*-test between groups in the row with *P* value shown in last column. Comparison only performed when repeat measured ANOVA showed $p < 0.001$ . *P* value in columns for comparison between vehicle and VC108 using unpaired *t*-test.

Figure 7 illustrates examples of IRV and INV from a Group 3A animals receiving either vehicle (panel A) or VC108 (0.5 mg/kg, panel B) 5 min prior to reperfusion The INV measured 60 min after reperfusion is significantly smaller in the VC108 versus vehicle treated animal. Panel C depicts the aggregate results from Group 3A animals, where INV normalized to the IRV is significantly smaller with VC108 compared to vehicle treated animals. Like Group 1A animals, pO_2_ during coronary occlusion does not have a positive influence on INV being higher in vehicle despite larger INV compared to VC108 treated animals.

**Figure 7:**
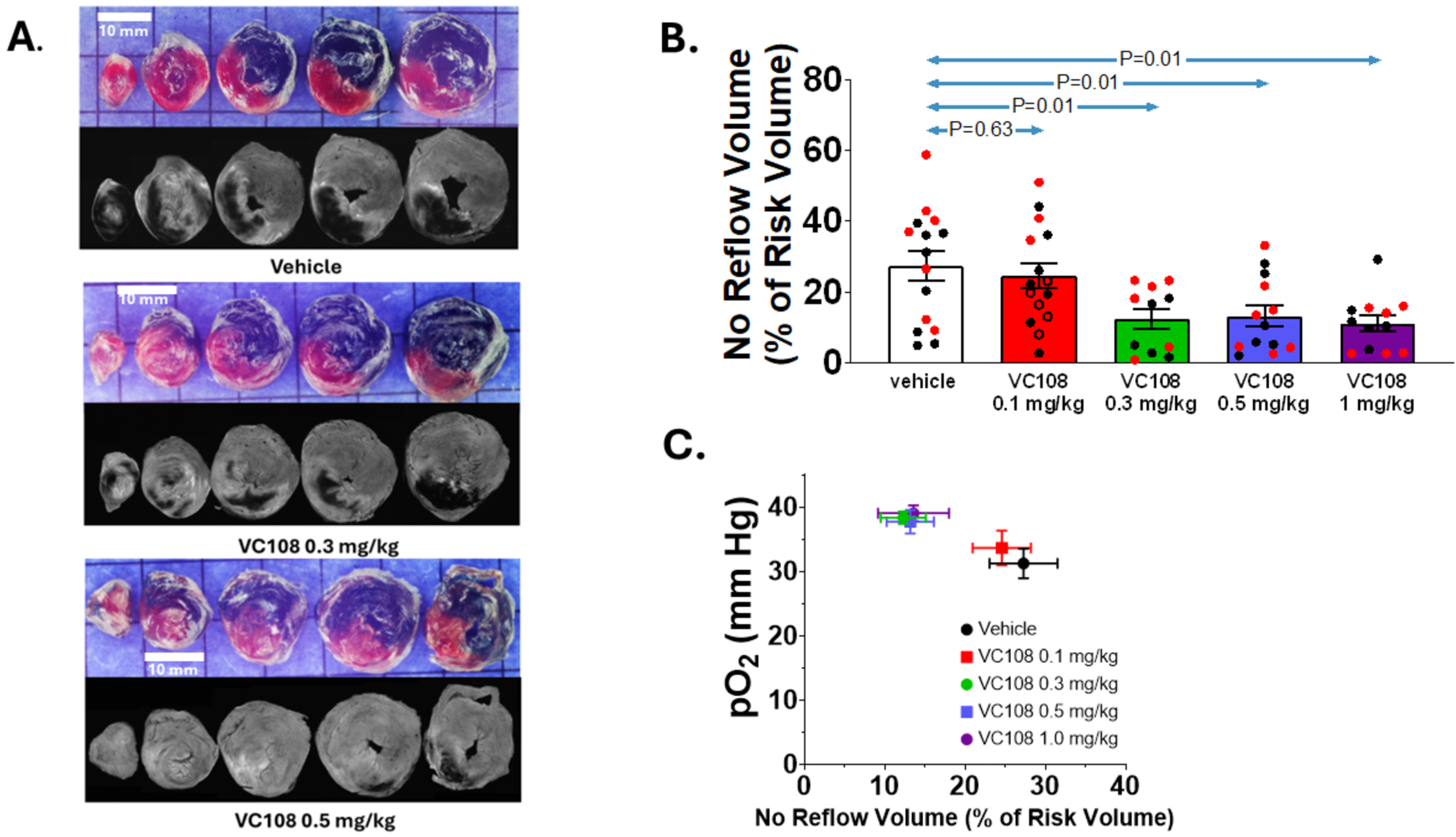
Results from Group 3A animals. Examples of INV (pale tissue) within IRV (tissue not stained blue) from a Group 3A animal receiving either (A) vehicle or (B) 0.5 mg/kg of VC108 5 min prior to reperfusion (scale = 10 mm). Aggregate data (C) show significantly smaller INV/IRV in VC108 treated (n=14) versus vehicle treated (n=15) animals. Males are depicted in black and females in red. No effect of tissue pO_2_ on the INV/IRV ratio is seen (D). Differences between mean were analyzed using unpaired t-test.

Table 5A shows the effect of different doses of VC108 given 5 min before reperfusion in Group 3B animals on heart rate, mean aortic pressure, WT and pO_2_ measured 35 min later during reperfusion. There was no difference in heart rate and mean aortic pressure between the stages for vehicle or any dose of VC108. However, heart rate at baseline and during coronary occlusion tended to be lower prior to administering 0.1 mg/kg of VC108 compared to before administering vehicle (Table 5B). While mean aortic pressure declined for vehicle and all VC108 doses during coronary occlusion, this decline reached statistical significance only in animals that would receive 0.1 mg/kg of VC108 prior to reperfusion. WT declined precipitously during coronary occlusion in all animals and then only partially recovered during reperfusion. WT data were not acquired at 0.1 mg/kg of VC108. pO_2_ declined significantly in all animals during coronary occlusion and recovered only partially during reperfusion. Compared to vehicle, pO_2_ was significantly higher during coronary occlusion in animals receiving 0.5 mg/kg of VC108 prior to reperfusion and during reperfusion at all doses of VC108 except for 0.1 mg/kg (Table 5B).

**Table 5A.** Results (mean ± 1SEM) From 66 Group 3B Animals Receiving different doses of VC108/Vehicle 5 min Before Reperfusion.

| **Drug Dose** | **Stage** | **Heart Rate**<br>**(bpm)** | **Aortic**<br>**Pressure**<br>**(mmHg)** | **% Wall**<br>**Thickening** | **pO<sub>2</sub> (mmHg)** |
| --- | --- | --- | --- | --- | --- |
| **Vehicle** | Baseline | 333± 19* | 86 ± 4* | 40.1 ± 0.3* | 41.4 ± 2.0* |
| (n=15) | Occlusion | 342 ± 10* | 78± 3* | 17.4 ± 2.6* | 11.2 ± 1.5* |
| | Reperfusion | 315 ± 10 | 75 ± 5 | 29.8 ± 1.8 | 31.3 ± 2.3 |
| **P value** |  | 0.6798 | 0.0873 | <0.0001 | <0.0001 |
| **0.1 mg/kg**<br>**VC108** | Baseline | 289 ± 10* | 92 ± 2* | | 42.9 ± 1.5* |
| (n=15) | Occlusion | 303 ± 9* | 79 ± 3* | | 4.6 ± 0.9* |
| | Reperfusion | 291 ± 14 | 73 ± 3 | | 33.6 ± 2.5 |
| **P value** |  | 0.3031 | 0.0006 |  | <0.0001 |
| **0.3 mg/kg**<br>**VC108** | Baseline | 340± 11* | 94 ± 4* | 41.4 ± 0.3* | 42.3 ± 1.3* |
| (n=11) | Occlusion | 334 ± 7* | 81 ± 5* | 19.7 ± 2.4* | 14.7 ± 1.7* |
| | Reperfusion | 313 ± 13 | 80 ± 2 | 28.4 ± 2.8 | 38.4 ± 0.9 |
| **P value** |  | 0.6089 | 0.0668 | <0.0001 | <0.0001 |
| **0.5 mg/kg**<br>**VC108** | Baseline | 319 ± 14* | 77 ± 5* | 40.2 ± 0.3* | 42.9 ± 2.0* |
| (n=13) | Occlusion | 340 ± 8* | 70 ± 5* | 18.1 ± 2.0* | 17.6 ± 2.5* |
| | Reperfusion | 315 ± 9 | 69 ± 5 | 30.3 ± 1.6 | 37.8 ± 1.8 |
| **P value** |  | 0.2155 | 0.3334 | <0.0001 | <0.0001 |
| **1 mg/kg**<br>**VC108** | Baseline | 325 ± 16* | 84± 3* | 41.0 ± 0.8* | 46.1 ± 1.9* |
| (n=12) | Occlusion | 343 ± 11* | 77 ± 3* | 13.6 ± 2.0* | 15.1 ± 1.9* |
| | Reperfusion | 286 ± 24 | 74 ± 3 | 31.3 ± 2.4 | 39.1 ± 1.2 |
| **P value** |  | 0.3759 | 0.0891 | <0.0001 | <0.0001 |

\*Comparison using unpaired *t*-test between groups in the columns with *P* value shown below. Wall thickening was not performed in animals receiving 0.1 mg/kg of VC108.

**Table 5B.** *P*-values in comparison to Vehicle.

|  | **stage** | **0.1 mg/kg** | **0.3 mg/kg** | **0.5 mg/kg** | **1 mg/kg** |
| --- | --- | --- | --- | --- | --- |
| **Heart Rate**<br>**(bpm)** | Baseline | 0.0543 | 0.7838 | 0.5690 | 0.7610 |
|  | Occlusion | 0.0090 | 0.5306 | 0.8313 | 0.9808 |
|  | Reperfusion | 0.1645 | 0.9277 | 0.9986 | 0.2237 |
| **Aortic**<br>**Pressure**<br>**(mmHg)** | Baseline | 0.1763 | 0.1871 | 0.1083 | 0.6960 |
|  | Occlusion | 0.8680 | 0.6353 | 0.1252 | 0.7086 |
|  | Reperfusion | 0.8108 | 0.3925 | 0.4310 | 0.9839 |
| **pO<sub>2</sub>**<br>**(mmHg)** | Baseline | 0.5539 | 0.7306 | 0.5928 | 0.1066 |
|  | Occlusion | 0.0008 | 0.1414 | 0.0337 | 0.1199 |
|  | Reperfusion | 0.4964 | 0.0216 | 0.0418 | 0.0115 |

Figure 8A shows IRV and NRV from 3 Group 3B animals given vehicle, 0.3 mg/kg and 0.5 mg/kg of VC108 5 min prior to reperfusion. NRV decreased with increasing doses of VC108 compared to vehicle. Figure 8B depicts the aggregate results of NRV from all Group 3B animals that received vehicle, and 4 different doses of VC108 prior to reperfusion. The NRV/IRV ratio at the lowest dose (0.1 mg/kg), which results in approximately 80% receptor occupancy, did not differ significantly from NRV/IRV ratio when vehicle was given. All other doses caused a significant (>50%) reduction in NRV/IRV compared to vehicle with *p* values increasing with increasing doses of VC108. However, when NRV/IRV was compared between 0.3, 0.5, and 1.0 mg/kg doses, no differences were noted. When NRV was multiplied by the ratio of mean pixel intensity in the no reflow zone versus the normal myocardium, the results were quite similar.

**Figure 8:**
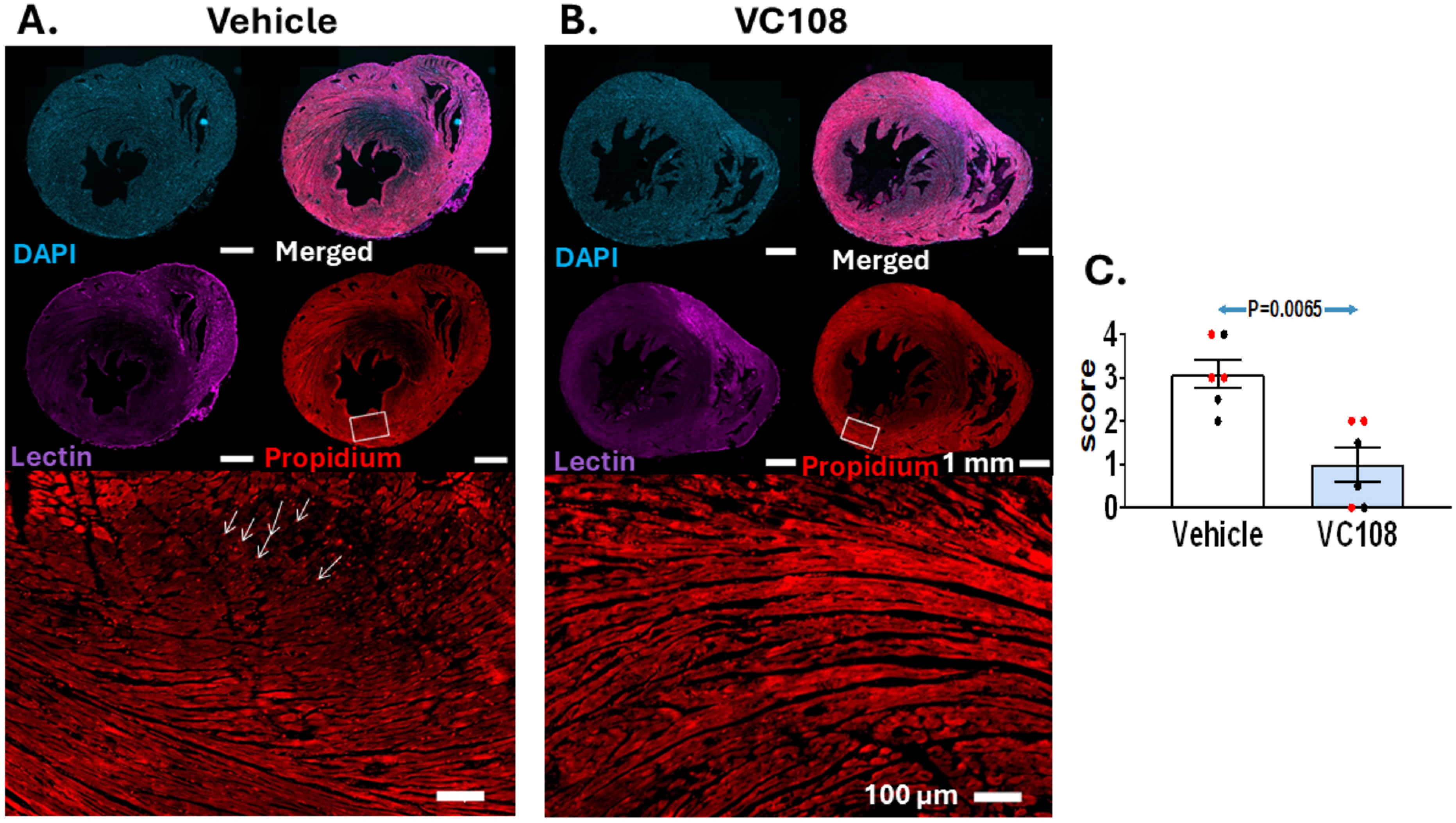
Results from Group 3B animals. Examples of NRV from 3 Group 3B animals receiving either vehicle (top panel) or 2 different doses (0.3 mg/kg, middle panel) and 0.5 mg/kg (lower panel) of VC108 5 min prior to reperfusion (A). Compared to vehicle, animals receiving VC108 demonstrated smaller NRV with the one receiving 0.5 mg/kg exhibiting the smallest NRV despite similar IRV in all animals (scale = 10 mm). Aggregate results from Group 3B animals (B) receiving vehicle (n=15) versus different doses of VC108 5 min prior to reperfusion. NRV/IRV ratio (A) is smaller for the 3 higher VC108 doses: 0.1mg/kg (n=15); 0.3mg/kg (n=11); 0.5 mg/kg (n=13); and 1.0 mg/kg (n=12). Males are depicted in black and females in red. Tissue pO_2_ at time of reperfusion (B) had a positive effect on the NRV/IRV ratio (C). Differences between mean were analyzed using unpaired t-test.

Figure 8C illustrates the positive effect of tissue pO_2_ during reperfusion on the NRV/IRV ratio. NRV/IRV was highest with the lowest pO_2_ value when vehicle was given. With 0.1 mg/kg VC108, pO_2_ was marginally higher and NRV/IRV was slightly lower. pO_2_ and NRV/IRV were superimposable with higher VC108 doses, where pO_2_ values were higher and NRV/IRV were lower.

To determine whether the reduction in INV with VC108 was influenced by its direct effect on cardiomyocytes, 12 additional animals underwent 1 h or coronary occlusion followed by 1 h of reperfusion (Group 3C) and 5 min before reperfusion were given either vehicle (n=6) or VC108 (n=6). At the end of the experiment, heart was removed for histopathology for necrosis, apoptosis, and ferroptosis.

Propidium iodide was used for assessment of necrosis. Figure 9 shows examples from vehicle treated (panel A) and VC108 treated (panel B) animals. Short-axis slices are shown with staining for DAPI, Lectin, and propidium iodide. The lectin and propidium iodide staining was performed in vivo and DAPI staining was performed ex vivo. The lectin image shows the risk area during coronary occlusion. Under high powered microscopy nuclei staining with propidium iodide are marked by arrows in the vehicle treated animal. Semi-quantitative analysis revealed marked reduction in propidium iodide staining in VC108 compared to vehicle treated animals (panel C).

**Figure 9:**
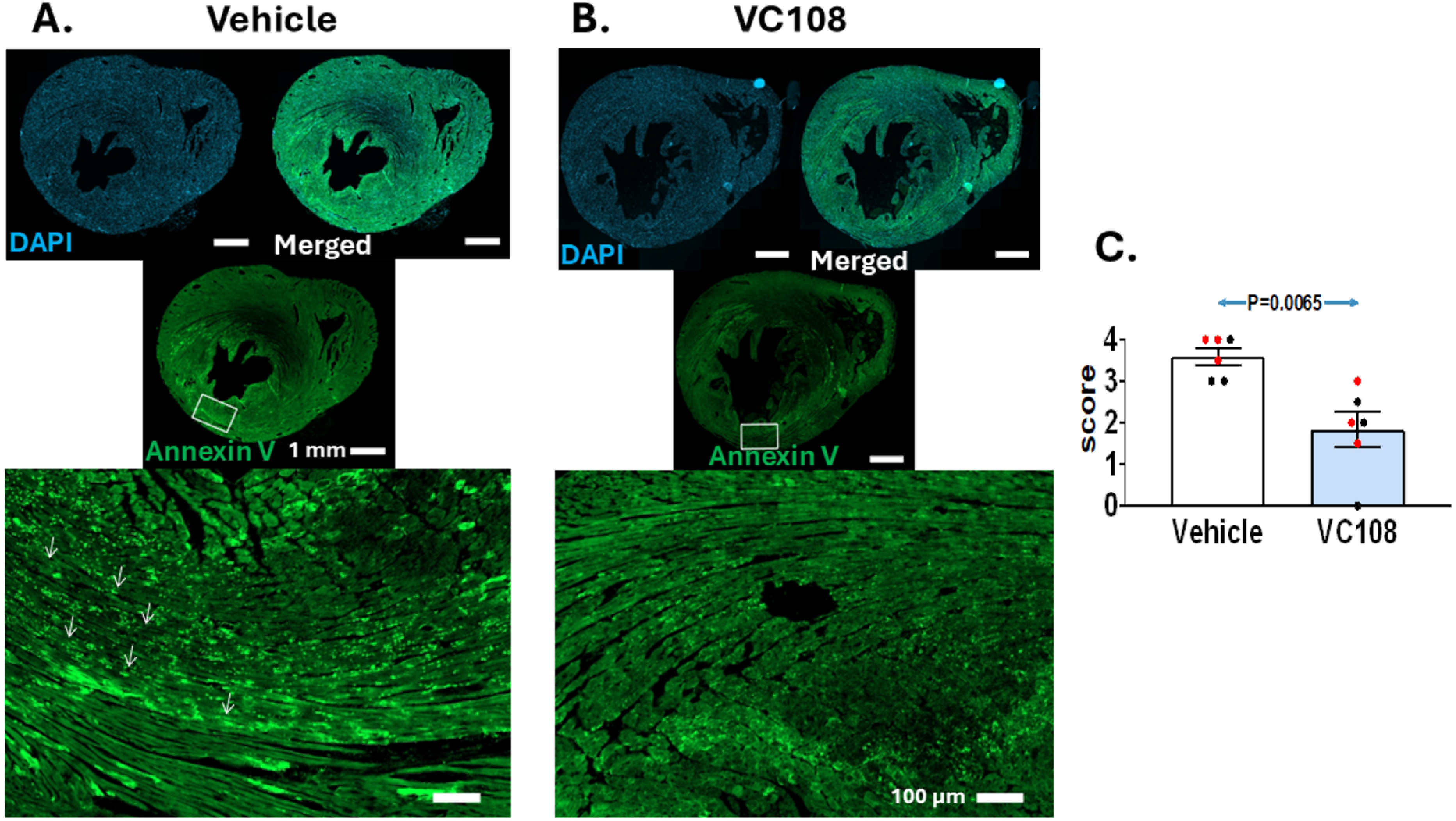
Assessment of necrosis by propidium iodide staining. Short-axis slices from vehicle treated (panel A) and VC108 treated (panel B) are shown with staining for DAPI, Lectin, and propidium iodide. The lectin image shows the risk area during coronary occlusion. Under high powered microscopy nuclei staining with propidium iodide are marked by arrows in the vehicle treated animal. Semi-quantitative analysis revealed marked reduction in propidium iodide staining in VC108 compared to vehicle treated animals (panel C). Males are depicted in black and females in red.

Annexin V was used for assessment of apoptosis. Examples of short-axis slices from vehicle treated (panel A) and VC108 treated (panel B) animals are shown stained with DAPI and Annexin V, along with merged image in Figure 10. On high powered microscopy annexin V entry into cells is seen in the vehicle treated and not the VC108 treated animals, indicating greater apoptosis in the former. Semi-quantitative analysis revealed considerable reduction in annexin V staining in VC108 compared to vehicle treated animals (panel C) confirming more apoptosis in the vehicle compared to VC108 treated animals.

**Figure 10:**
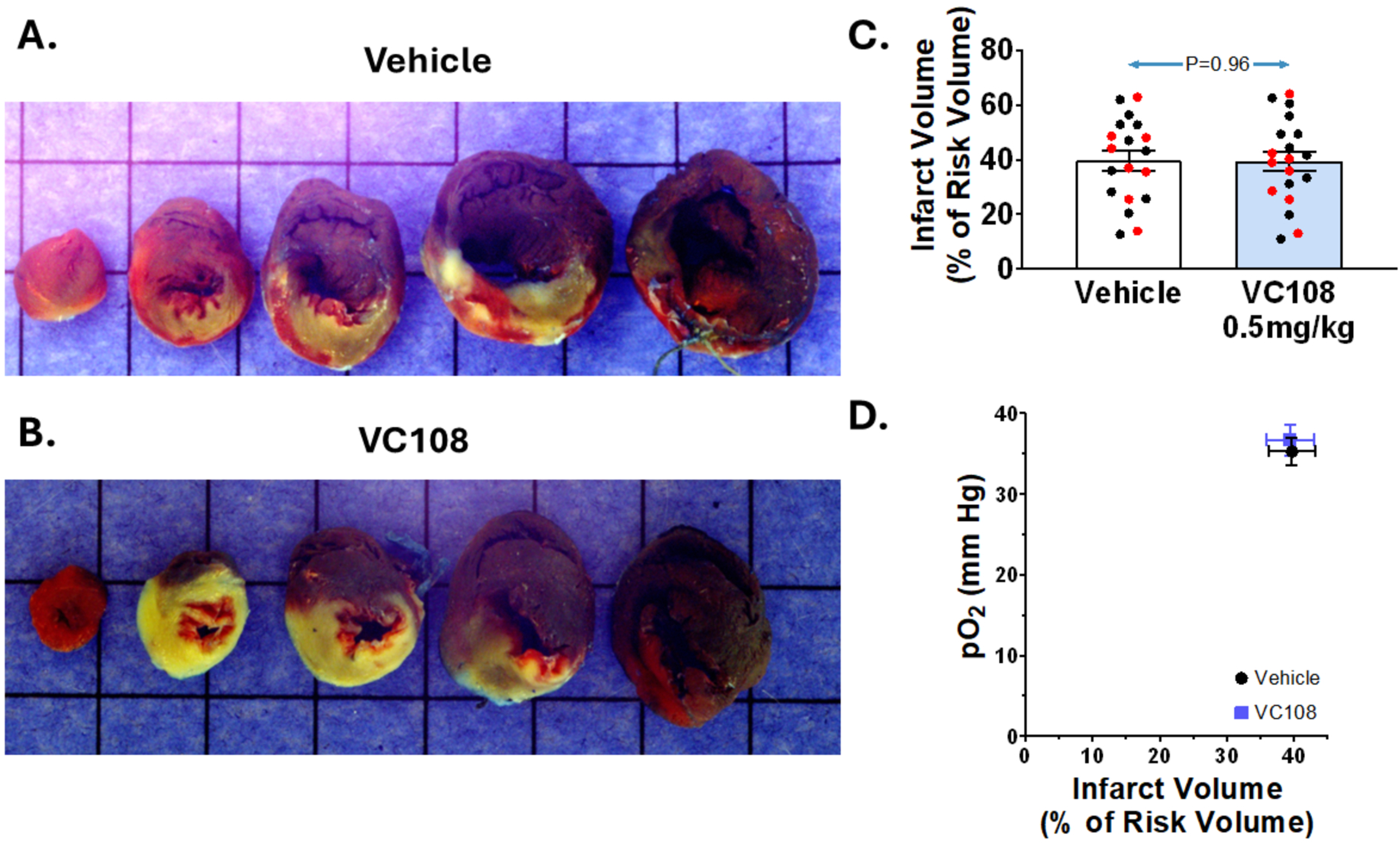
Assessment of apoptosis by Annexin V staining. Examples of short-axis slices from vehicle treated (panel A) and VC108 treated (panel B) animals are shown stained with DAPI and Annexin V, along with merged image. On high powered microscopy annexin V entry into cells is seen in the vehicle treated and not the VC108 treated animals, indicating greater apoptosis in the former. Semi-quantitative analysis revealed considerable reduction in annexin V staining in VC108 compared to vehicle treated animals (panel C) confirming more apoptosis in the vehicle compared to VC108 treated animals. Males are depicted in black and females in red.

Supplemental Figure 1 shows results of ferroptosis. We found no difference between vehicle and VC108 for iron staining of infarcted tissue.

### Group 4 animals

Table 6 depicts results from the Group 4 animals who received 0.5 mg/kg or VC108 30 min after reperfusion and in which INV and NRV were measured 60 min after reperfusion. Heart rate increased after coronary occlusion in both vehicle and VC108 treated animals. Mean aortic pressure did not change between the stages and were not different at any stage between vehicle and VC108. pO_2_ was significantly higher in VC 108 treated animals prior to coronary occlusion and declined precipitously during coronary occlusion to the same extent as vehicle treated animals. After reperfusion, pO2 increased equally in the vehicle and VC108 treated animals.

**Table 6.** Results (mean ± 1SEM. ) **from Group 4 Animals Receiving 0.5mg/kg VC108 (n=38) or vehicle (n=35) 30 min After Reperfusion**

| **Variable** | **Pre-**<br>**Occlusion** | **Occlusion** | **Reperfusion** | ***P* value** |
| --- | --- | --- | --- | --- |
| **Heart Rate (bpm)** |  |  |  |  |
| **Vehicle** | *319 ± 4 | *347 ± 6 | 325 ± 6 | 0.0003 |
| **VC108** | *310 ± 4 | *338 ± 8 | 328 ± 5 | 0.0003 |
| ***P* value** | 0.1301 | 0.2747 | 0.7081 |  |
| **Aortic Pressure**<br>**(mm Hg)** |  |  |  |  |
| **Vehicle** | *95 ± 2 | *91 ± 2 | 73 ± 2 | 0.1263 |
| **VC108** | *97 ± 2 | *91 ± 2 | 74 ± 2 | 0.0553 |
| ***P* value** | 0.6089 | 0.7858 | 0.7149 |  |
| **pO<sub>2</sub> (mm Hg)** |  |  |  |  |
| **Vehicle** | *38.6 ± 1.3 | *5.7 ± 0.4 | 36.8 ± 1.3 | <0.0001 |
| **VC108** | *43.0 ± 1.2 | *5.6 ± 0.6 | 38.4 ± 1.2 | <0.0001 |
| ***P* value** | 0.0151 | 0.9274 | 0.3672 |  |

\*Comparison using paired *t*-test between pre- and post-treatment during coronary occlusion with *P* value shown in last column. *P* value in columns for comparison between vehicle and VC108 using unpaired *t*-test. Wall Thickening was not performed for these animals.

Figure 11 illustrates IRV and INV from a Group 4A animal receiving vehicle (panel A) and 0.5 mg/kg of VC108 (panel B) 30 min after reperfusion. The INV is similar between drug and vehicle treated animals. Aggregate results from Group 4A animals show that INV/IRV is similar between vehicle and VC108 treated animals (panel C). Tissue pO_2_ during reperfusion measured after vehicle/drug administration had no relation to INV/IRV ratio (panel D).

**Figure 11:**
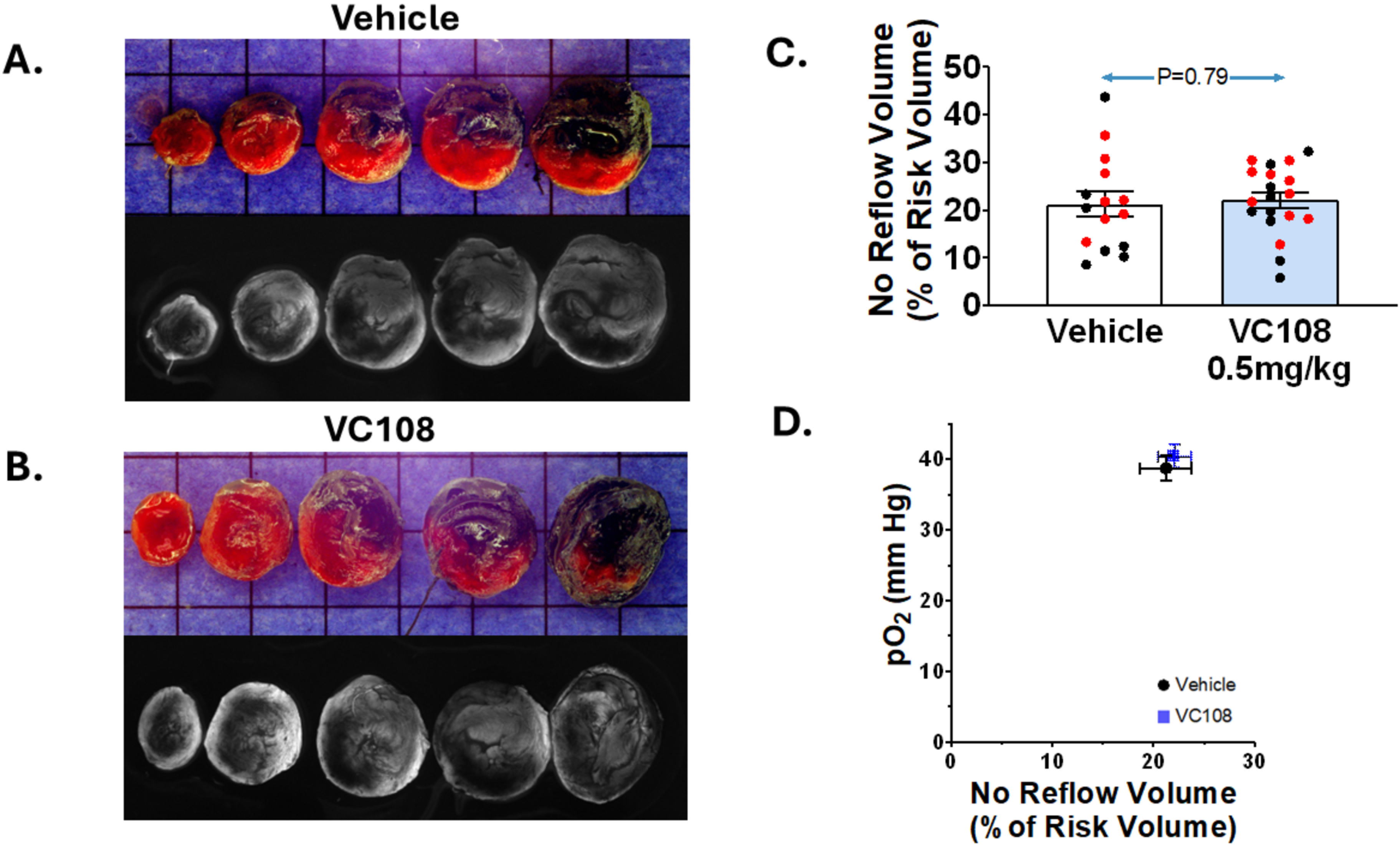
Results from Group 4A animals. Examples of INV (pale tissue) within IRV (tissue not stained blue) from a Group 4A animal receiving either (A) vehicle or (B) 0.5 mg/kg of VC108 30 min after reperfusion (scale=10 mm). Aggregate data (C) show similar INV/IRV in VC108 treated (n=19) versus vehicle treated (n=19) animals. Males are depicted in black and females in red No effect of tissue pO_2_ on the INV/IRV ratio is seen (D). Differences between mean were analyzed using unpaired t-test.

Figure 12 illustrates IRV and NRV from a Group 4B animal receiving vehicle (panel A) and 0.5 mg/kg of VC108 (panel B) 5 min prior to reperfusion. The NRV is similar between drug and vehicle treated animals. Aggregate results from Group 4B animals show that NRV/IRV is similar between VC108 and vehicle treated animals (panel C). Tissue pO_2_ during reperfusion measured after vehicle/drug administration had no relation to NRV/IRV ratio (panel D).

**Figure 12:**
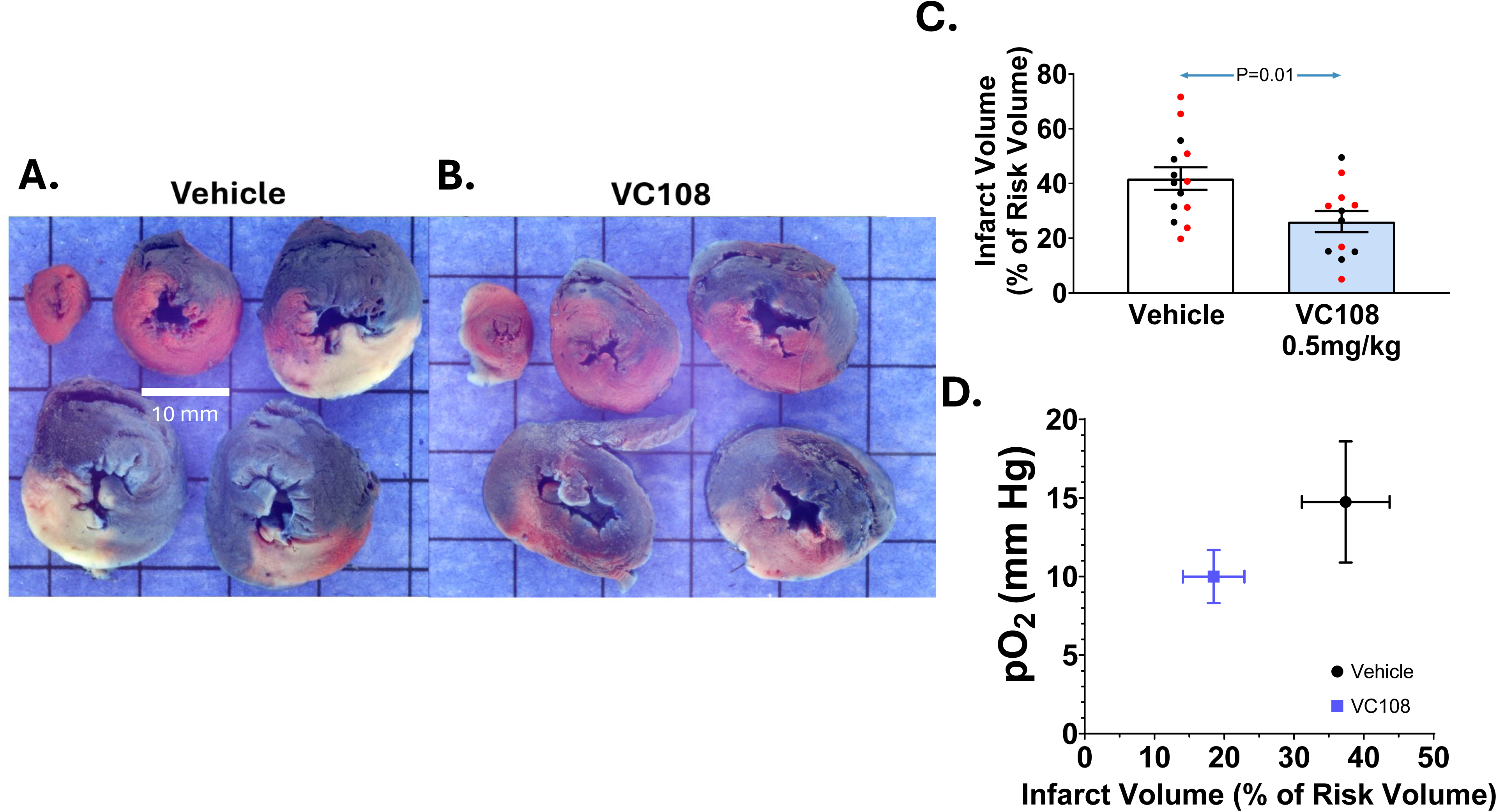
Results from Group 4B animals. Examples of NRV from a Group 4B animal receiving either vehicle (top panel) receiving either (A) vehicle or (B) 0.5 mg/kg of VC108 30 min after reperfusion (scale=10 mm). Aggregate data (C) show similar NRV/IRV in VC108 treated (n=19) versus vehicle treated (n=15) animals. Males are depicted in black and females in red. No effect of tissue pO_2_ on the NRV /IRV ratio is seen (D). Differences between mean were analyzed using unpaired t-test.

### Group 5 animals

Table 7A lists the results of the total and free plasma as well as tissue concentrations of VC108 at 3 different doses in 6 animals each where samples were obtained 45 min after coronary occlusion without performing reperfusion. Although the maximal dose of the drug was 10 times higher than the lowest dose (5 mg/kg vs 0.5 mg/kg) the plasma concentration increased by only 1.4 times, while the tissue concentrations both in the risk area and normal myocardium increased by 3-fold and 3.8-fold, respectively. The risk area/plasma ratio increased by 2-fold at the highest compared to the lowest dose. Interestingly, the ratio of drug concentration in the risk area versus normal myocardium was close to 1 indicating good penetration of drug in the ischemic bed. The in vitro IC50 of VC108 is 64.88 nM. The free plasma concentration of the drug exceeds the IC50 at all doses administered.

**Table 7A.** Total (ng/mL) and Free (nM) Plasma and Tissue Levels (ng/G) of VC108 From Group 5 Animals 45 min After Receiving Intravenous Doses of Drug Administered 15 min After Coronary Occlusion (mean±1SEM)

| **Dose (mg/kg)** | **Plasma Concentration** | **Myocardial risk Area Concentration** | **Normal Myocardial Concentration** | **Risk Area/ Normal Myocardium Ratio** | **Risk area/ Plasma ratio** | **Normal Myocardium/ Plasma ratio** |
| --- | --- | --- | --- | --- | --- | --- |
| 0.5 (n=6) | T: 384.9 ± 55.1<br>F: 273.2 ± 39.1 (4.2) | 259.6 ± 31.2 | 232.1 ± 25.9 | 1.1 ± 0.18 | 0.68 ± 0.44 | 0.60 ± 0.22 |
| 2.5 (n=6) | T: 511.7 ± 60.1<br>F: 363.2 ± 42.7 (5.6) | 576.3 ± 69.7 | 603.4 ± 72.8 | 1.1 ± 0.13 | 1.14 ± 0.10 | 1.13 ± 0.18 |
| 5.0 (n=6) | T: 556.8 ± 81.9<br>F: 395.2 ± 58.1 (6.1) | 801.5 ± 154.6 | 873.6 ± 177.6 | 0.99 ± 0.09 | 1.47 ± 0.23 | 1.35 ± 0.23 |

### Group 6 Animals

Table 7B illustrates results of blood levels of VC108 at 5, 10, 15, 30, and 60 min after injection of 3 different doses of VC108 5 min prior to reperfusion. Tissue drug levels from ischemic and normal myocardium were measured 60 min after injection. Plasma drug levels declined over time for all doses with tissue levels at 60 min being slightly higher than plasma levels at that time point for both risk area and normal myocardium. Unlike the Group 5 animals, however, the plasma levels at all time-points increased linearly with dose administered. As in Groups 5 animals, the tissue level of drug was equivalent in the normal and ischemic tissue at this time point. The free plasma concentration of the drug exceeds the IC50 at all time points at 1 mg/kg dose but not at 0.1 mg/kg. For 0.3 mg/kg it is adequate early after drug administration.

**Table 7B.** Total (ng/mL) and Free (nM) Plasma and Tissue Levels (ng/G) of VC108 From Group 6 Animals Receiving Intravenous Doses of Drug 5 min Prior to reperfusion (mean±1SEM)

| **Dose (mg/kg)** | **Plasma Concentration 5 min** | **Plasma Concentration 10 min** | **Plasma Concentration 15 min** | **Plasma Concentration 30 min** | **Plasma Concentration 60 min** | **Myocardial risk area concentration 60 min** | **Normal myocardial concentration 60 min** |
| --- | --- | --- | --- | --- | --- | --- | --- |
| 0.1 (n=3) | T: 53.3 ± 9.1<br>F: 37.8 ± 6.5<br>(0.6) | T: 29.5 ± 0.7<br>F: 20.9 ± 0.5<br>(0.32) | T: 24.1 ± 1.5<br>F: 17.1 ± 1.0<br>(0.3) | T: 15.7 ± 4.0<br>F: 11.1 ± 2.8<br>(0.8) | T: 7.23 ± 0.5<br>F: 5.1 ± 0.3<br>(0.1) | 8.72 ± 1.4 | 9.01 ± 1.3 |
| 0.3 (n=3) | T: 82.9 ± 19.2<br>F: 58.8 ± 16.6<br>(0.9) | T: 48.7 ± 8.2<br>F: 34.6 ± 5.8<br>(0.5) | T: 38.7 ± 2.9<br>F: 27.5 ± 2.9<br>(0.4) | T: 22.6 ± 9.5<br>F: 16.0 ± 6.7<br>(0.2) | T: 9.82 ± 1.8<br>F: 7.0 ± 1.3<br>(0.1) | 13.0 ± 3.6 | 15.4 ± 5.8 |
| 1.0 (n=3) | T: 405.0 ± 38.9<br>F: 287.5 ± 27.6<br>(4.4) | T: 263.8 ± 27.1<br>F: 187.2 ± 13.2<br>(2.9) | T: 261.0 ± 40.9<br>F: 185.3 ± 29.0<br>(2.8) | T: 263.0 ± 63.6<br>F: 186.7 ± 45.0<br>(2.9) | T: 83.5 ± 20.6<br>F: 59.0 ± 14.6<br>(0.9) | 109.0 ± 19.3 | 123.0 ± 41.6 |

T= total plasma concentration (ng/mL); F=Free plasma concentration (nM); ( ) Free plasma/IC50 ratio

### Pharmacokinetics of VC108

Following a single IV bolus injection of 1 mg/kg VC108 was quantifiable in plasma up to at least 1 h after dosing (T_last_). Supplemental Figures 2 and 3 depict time versus plasma concentration in linear and logarithmic scales, respectively. Time after injection when maximal concentration was achieved (T_max_) was 0.083 hours. The mean maximal concentration (C_max_) was 797 ng/mL and mean area under the plasma concentration time curve from the time of dosing to the last measurable concentration (AUC_last_) was 320 ng·h/mL. The mean calculated half-life was 0.255 hours (range 0.219 and 0.284 hours). VC108 showed a moderate mean plasma clearance of 3030 mL/h/kg and a moderate volume of distribution (V_ss_) of 1040 mL/kg (higher than total body water volume).

Following single and repeated intravenous administration, systemic exposure increased with increasing dose of VC108 with no major deviation from proportionality for each dose in either male or female rats, except AUC_last_ in females, which increased with a tendency to supra-proportionality. Females were more exposed than males due to the slower elimination of VC108 characterized by a lower clearance and higher terminal half-life. No notable time dependent differences in systemic exposure were observed between single and repeated administrations. A summary of parameters is shown below in Supplemental Table 2.

Of note, the doses of VC108 used in the AMI studies were significantly lower than the doses used in the pharmacokinetic studies other than the 3 male rats who received 1 mg/kg bolus.

## Discussion

The new information from this study is that GPR39 is present not only in rat heart VSMCs and pericytes as noted previously in mice^1.2^, but also in cardiomyocytes. Specific pharmacological inhibition of GPR39 by VC108, a drug that has completed pre-clinical evaluation for new investigational drug approval as a treatment for AMI, resulted in marked reduction in both INV and NRV when given prior to coronary occlusion or just prior to reperfusion in AMI. It had no effect when administered 30 min after reperfusion. VC108 also caused a marked decrease in INV when administered during coronary occlusion. Interestingly, higher tissue pO_2_ with VC108 during ischemia indicating enhanced perfusion, was always associated with less NRV no matter when VC108 was injected: before and during coronary occlusion or prior to reperfusion. In comparison, increased tissue pO_2_ was associated with INV reduction only when VC108 was administered during coronary occlusion. Further exploration revealed that VC108 resulted in less necrosis, and apoptosis compared to vehicle, implying that it directly affected cardiomyocytes independent of its effect on tissue pO_2._

15-HETE is the endogenous agonist for GPR39. It is metabolized from arachidonic acid predominantly by 15-lipoxygenase (15-LO) but also by cyclogeneses and cytochrome 450.^21,22^ 15-HETE levels are elevated in human ischemic myocardium.^23^ Cardiomyocytes and endothelial cells^24^ exposed to hypoxia also express higher levels of 15-HETE. We showed that 15-HETE increases intracellular Ca^++^ resulting in VSMC and pericyte contraction.^1,2^ Our current results show that GPR39 is also present in cardiomyocytes, which may facilitate intracellular Ca^++^ release and overload during ischemia that is associated with mitochondrial damage and myocardial dysfunction. Here we show that, compared to vehicle, VC108 results in less necrosis and apoptosis within the ischemic zone. Selective pharmacological inhibition of GPR39 may thus confer protection to ischemic myocardium through more than enhanced tissue oxygenation caused by vasodilation.

### Direct Effects of 15-HETE on Cardiomyocytes

Apart from causing cardiomyocyte contraction, Ca^++^ also acts as a second messenger that activates protein kinase C (PKC), which hydrolyses phosphatidylinositol 4,5-bisphosphate into membrane bound diacylglycerol (DAG) and cytosolic inositol 1,4,5-trisphosphate (IP3). The latter acts on IP3 receptors in the endoplasmic reticulum to release more Ca^++^. DAG also activates PKC that in turn causes phosphorylation of various proteins associated with apoptosis and necrosis.^25^ Inhibition of PKC-δ during reperfusion can protect against ischemic damage by preventing its translocation to the mitochondria. Thus, GPR39 blockade could be beneficial in myocardial ischemia by inhibiting PKC activation. Our results indeed show reduced necrosis and apoptosis following pharmacological inhibition of VC108.

Free Ca^++^ also forms a complex with calmodulin, activating calcium/calmodulin dependent protein kinases (CAMK). Activation of CAMKII is associated with inflammation and cell death in myocardial ischemia and reperfusion.^26^ GPR39 inhibition could, hence, reduce CAMKII activation and prevent cell death as well as inflammation. In our current study we did not assess inflammation.

We demonstrated that 15-HETE activated extracellular regulated kinase (ERK)-mitogen associated protein kinase (MAPK) pathway by stimulating Gα_q_.^1^ In ischemia/reperfusion this pathway regulates ferroptosis, a form of cell death characterized by iron-dependent lipid peroxidation.^27^ Whereas we were unable to demonstrate reduced presence of iron in VC108 treated animals, we did not measure glutathione and glutathione peroxidase 4. Thus, our assessment of ferroptosis may not have been as comprehensive. MAPK signaling also increases apoptosis,^28^ the reduction of which was seen with VC108. In addition, the macrophage associated inflammasome leading to fibrosis is also activated by 15-HETE.^29^ Thus GPR39 inhibition could mitigate these short and intermediate term harmful effects of 15-HETE.

There are no reports implicating 15-HETE-GPR39 signaling through Gα_s_ or Gα_12/13_. However, supraphysiological levels of Zn^++^ as well as synthetic agonists have been shown to stimulate Gα_s_ leading to cyclic adenosine monophosphate (cAMP) response element-mediated transcription induction^30^, which along with protein kinase A, can lead to altered expression of genes associated with apoptosis and necrosis, both of which were reduced by VC108 in our study. Stimulation of Gα_12/13_ induces serum response element (SRE)-mediated transcription that affects cell survival.^31^ GPR39 stimulation could also cause other downstream signaling that may be detrimental to cells.^32–34^

### Vasodilation Secondary to GPR39 blockade

In a previous study we showed that genetic deletion of GPR39 reduced NRV in a mouse model of AMI that was associated with increased capillary density and diameter within the risk area compared to littermate wild-type mice, suggesting a role of pericytes in contributing to myocardial ischemia.^2^ Pericytes were previously implicated in reducing capillary perfusion in myocardial ischemia, where administration of adenosine relaxed the pericytes and increased perfusion.^35^ We reported that when perfusion pressure falls distal to a stenosis during exogenous hyperemia, pericytes contract causing capillary constriction.^36^ During myocardial ischemia, arterioles (VSMCs) are fully dilated indicating that any increase in perfusion caused by an intervention has to relax pericytes since in this situation their contraction is the main factor limiting nutrient flow.^37^

Whereas the arterial effects of 15-HETE has been mostly studied in pulmonary hypertension, it has been shown to increase vasoconstriction in cerebral, coronary, and umbilical arteries as well.^38–40^ Apart from Ca^++^ induced VSMC and pericyte contraction, 15-HETE also inhibits K^+^ channels during cerebral hypoxia.^38^ Vasodilation requires VSMC hyperpolarization that is mediated through increased levels of intracellular K^+^ causing VSMC (and pericyte?) K^+^ channel opening. Hence, GPR39 blockade would result in more K+ channel opening, providing another mechanism for VSMC and pericyte relaxation and resulting vasodilation, leading to increased blood flow during myocardial ischemia.

### Blood and Tissue Concentrations of VC108

The plasma half-life of 1 mg/kg of VC108 administered as a bolus in rats is 0.25 h. Forty-five min after coronary occlusion, the concentration of drug within the ischemic bed was two-thirds of that in plasma at the 0.5 mg/kg dose that occupies >90% of GPR39 receptors in the rat. This was equivalent to the concentration in the normal myocardium implying there was enough perfusion within the ischemic tissue for the drug to accumulate over 45 min after it was administered. The free plasma concentration exceeded the IC50 at all doses, consistent with the biological effect seen in Group 2 animals. The non-linear increase in plasma concentration with higher doses probably indicates plasma protein saturation of drug and clearance of unbound drug.

There was linear relation between drug dose and total and free plasma concentration at all time points after reperfusion and the tissue concentration after 60 min of reperfusion was only slightly less in the erstwhile ischemic compared to the normal myocardium, and marginally higher than in plasma at that time indicating drug accumulation in tissue. The free plasma concentration of the drug always exceeded the IC50 at 1 mg/kg, but not at 0.1 mg/kg dose at all time points after reperfusion. These results are consistent with the biological effects seen in Group 3 animals.

### Critique of our methods

We administered VC108 or vehicle prior to and during coronary occlusion as well as just prior to and 30 min after reperfusion. The pre-occlusion administration was meant to demonstrate proof that VC108 has beneficial effects. Administration during coronary occlusion was to simulate a more clinically relevant situation. Finally, the pre-reperfusion administration was meant to mimic the situation where a patient presents to the emergency room for a possible percutaneous intervention for AMI. A single intravenous bolus was meant to facilitate logistically easy and timely intervention in the cardiac catheterization laboratory. We show that administering the drug 30 min after reperfusion has no beneficial effect. It would be unlikely that pericyte relaxation would have any additional benefits in the presence of reactive hyperemia at this stage that can last for hours or days.^41–43^ Further, by this time cardiomyocytes are likely irreversibly damaged in our AMI model, and drugs are unlikely to offer any salvage.

Our post-mortem methods for assessing NRV and INV are standard. Because these readouts cannot be made in the same animal, different sets of animals were used. It is difficult to measure tissue perfusion in rodents, hence, we decided to measure tissue O_2_ tension using a method we were familiar with.^14^ The ultimate reason to measure tissue perfusion is to define O_2_ delivery. O_2_ consumption was likely similar in all animals during ischemia or reperfusion and thus unlikely to independently affect pO_2_. The limitation of this method is that it reflects tissue pO_2_ only in the area where oxyphor is injected and cannot provide the spatial estimation of tissue pO_2_ within the entire IRV. However, since we occluded the proximal LC, ischemia always included the region where the oxyphor was injected.

### Translational Perspective

Historically, experimentally induced coronary (and other organ) no reflow after release of vessel occlusion was not associated with occurrence of microthrombi.^44–52^ However, when no reflow was reported in patients undergoing percutaneous coronary interventions (PCI)^53,54^ interventional cardiologists assumed that no reflow was caused by distal embolization of thrombus during PCI. Consequently, many studies were performed to demonstrate the benefit of pharmacologically or mechanically removing thrombus. Several large multicenter studies have now conclusively demonstrated that antithrombotic therapy or mechanical removal of thrombus does not improve coronary no reflow.^55–61^ No-reflow is independent of microthrombi. Microthrombi are present within the myocardium after PCI but dissolve because of aggressive antithrombotic therapy.^62,63^

During PCI (either just before or after opening the infarct-related artery) several vasodilators were given intracoronary in studies utilizing relatively small numbers of patients. Of these, neither nitroglycerin nor nitroprusside was shown to affect no reflow in larger studies.^64,65^ A K^+^ channel blocker, nicorandil that is not available in the USA, was shown to improve no-reflow^66^, which is not surprising given that pericytes have abundant K^+^ channels.^67^ Similarly, intracoronary Ca^++^ channel blockers were shown to reduce no reflow^68,69^ as was intracoronary epinephrine.^70,71^ However, requirement of selective intracoronary administration limits the use of these drugs.

Pericytes have abundant adenosine receptors.^67^ Topical administration of adenosine during experimental no reflow has been shown to markedly reduce no reflow.^35^ However, adenosine has a very short half-life and its intracoronary administration has produced questionable results.^64,72^ Attempts at giving it intravenously over several hours have shown favorable effects on LV function but have failed to reduce no reflow.^73,74^ When adenosine has shown reduction in no reflow it has been associated with adverse effects, such as hypotension and heart block.^75^ It is likely that achieving a high enough dose to inhibit pericyte contraction requires large intravenous doses that are poorly tolerated.

Unlike other vasodilators, VC108 has no systemic effects. It is a selective coronary vasodilator through its effects on both VSMCs and pericytes.^2,13^ As shown in this study, VC108 also has direct cardioprotective properties that have not been compared to other vasodilators. It is meant to be given as a single intravenous bolus while the patient is being prepped for PCI, which makes it logistically attractive. As shown in our study, the timing of drug administration also seems crucial.

### Conclusions

VC108, ready for application to the US Food and Drug Administration for Investigational Drug Approval as a single bolus injection for AMI, is very effective in reducing INV and NRV in a rat AMI model irrespective of timing of administration so long as it is given prior to reperfusion. It is also effective in reducing INV when administered during coronary occlusion. VC108 acts by blocking GPR39 for which the endogenous agonist is 15-HETE. This blockade results in vasodilation by relaxing pericytes and protection of cardiomyocytes by preventing other downstream harmful effects of GPR39 stimulation.

## Supporting information

https://doi.org/10.6084/m9.figshare.33154937

## Acronyms and Abbreviations

15-HETE: 15-hydroxyeicosatetraenoic acid
14,15-EET: 14,15-epoxyeicosatrienoic acid
AMI: acute myocardial infarction
cAMP: cyclic adenosine monophosphate
CMPK: calmodulin dependent protein kinase
DAG: diacylglycerol
ERK: extracellular regulated kinase
GPR39: G-protein coupled receptor 39
IHC: immunohistochemistry
IP3: inositol 1,4,5-trisphosphate
INV: infarct volume
IRV: ischemic risk volume
LC: left coronary
LV: left ventricular
MAPK: mitogen-activated protein kinase
NRV: no reflow volume
PCI: percutaneous coronary intervention
PKC: protein kinase C
qPCR: quantitative polymerase chain reaction
RNA: ribonucleic acid
TTC: 1% 2,3,5 triphenyl tetrazolium chloride
VSMC(s): vascular smooth muscle cell(s)
WT: wall thickening

## Supplemental Materials

Supplemental Tables 1 and 2

Supplemental Figures 1,2,3

Source data for supplemental material: 10.6084/m9.figshare.32948612

## Data Availability

Source data are available from the senior author upon request.

## Acknowledgements

The research reported in this publication used computational infrastructure supported by the Office of Research Infrastructure Programs, Office of the Director, of the National Institutes of Health under Award Number S10OD034224. The content is solely the responsibility of the authors and does not necessarily represent the official views of the National Institutes of Health.

We are thankful to the integrated genomic laboratory for use of qPCR equipment. It receives its support from the National Cancer Institute Center Support Grant P30CA069533 to the Knight Cancer Institute. We acknowledge expert technical assistance from the OHSU Advanced Light Microscopy Core (RRID:SCR_009961). We also acknowledge the use of Biorender.com to generate the central illustration.

## Grants

Supported, in part, by grants from the Garthe and Grace L. Brown Fund and the John E. and Robin Jaqua Fund at the Oregon Community Foundation and by the Ernest C. Swigert Endowment at the Oregon Health & Science University

## Disclosures

Oregon Health & Science University and Vasocardea, Inc. have a joint global patent (WO2021222858, and corresponding national filings) that encompasses drugs that inhibit GPR39. Vasocardea, Inc. has an additional patent filed globally (WO2023076219 and corresponding national filings) that encompasses VC108. Vasocardea, Inc. has exclusive rights to develop and commercialize drugs under all these patent applications.

Aptuit/Evotec has assigned all patent rights to Oregon Health & Science University and Vasocardea, Inc. Dr. Kaul is the founder and President of Vasocardea, Inc., a Delaware incorporated company located in Portland, Oregon.

## Disclaimer

The information in this manuscript does not represent the views of the Veterans Administration or the US government.

## Author Contributions

SK and CM (Methner) developed the concept and design for the studies and performed the experiments. CM (Methner) performed hemodynamic, pO_2_, western blot, and histopathology analysis as well as ICC. SK analyzed the post-mortem gross pathology. LL performed IHC and histopathology for necrosis, apoptosis, and ferroptosis. MP isolated rat pericytes and VSMCs. MK isolated the cardiomyocytes. AT performed qPCR. DEL performed the echocardiographic analysis. AC and FB performed medicinal chemistry for the development of VC108. AP performed the analysis in Groups 5 and 6 animals. The rat pharmacokinetic studies were performed by JF, CM (Marcheselli), and AP. Data curation and statistical analysis were performed by CM. All authors contributed to drafting the manuscript with SK drafting the final version.

## Notes

### Competing Interest Statement

Oregon Health & Science University and Vasocardea, Inc. have a joint patent application filed globally (WO2021222858 and corresponding national filings) that encompasses drugs that inhibit GPR39. Vasocardea, Inc. has an additional patent application filed globally (WO2023076219 and corresponding national filings) that encompasses VC108. Vasocardea, Inc. has exclusive rights to develop and commercialize drugs under all of these patent applications. Aptuit/Evotec has assigned all patent rights to Oregon Health & Science University and Vasocardea, Inc.
Dr. Kaul is the founder and President of Vasocardea, Inc., a Delaware incorporated company located in Portland, Oregon.
The information in this manuscript does not represent the views of the Veterans Administration or the US government.

### Summary of Updates

The paper has been revised as per reviewer's request. we added more animals and performed immunohistochemistry for ferroptosis, necrosis, and apoptosis.

