## Supplementary material for "Selective Pharmacological Blockade of GPR39 Markedly Reduces No Reflow and Infarct Volumes in a Rat Acute Myocardial Infarction When Administered Prior to Reperfusion": https://doi.org/10.6084/m9.figshare.33154937

**Supplemental Materials**

**Supplemental Table 1. In-vitro ADME Properties of VC108**

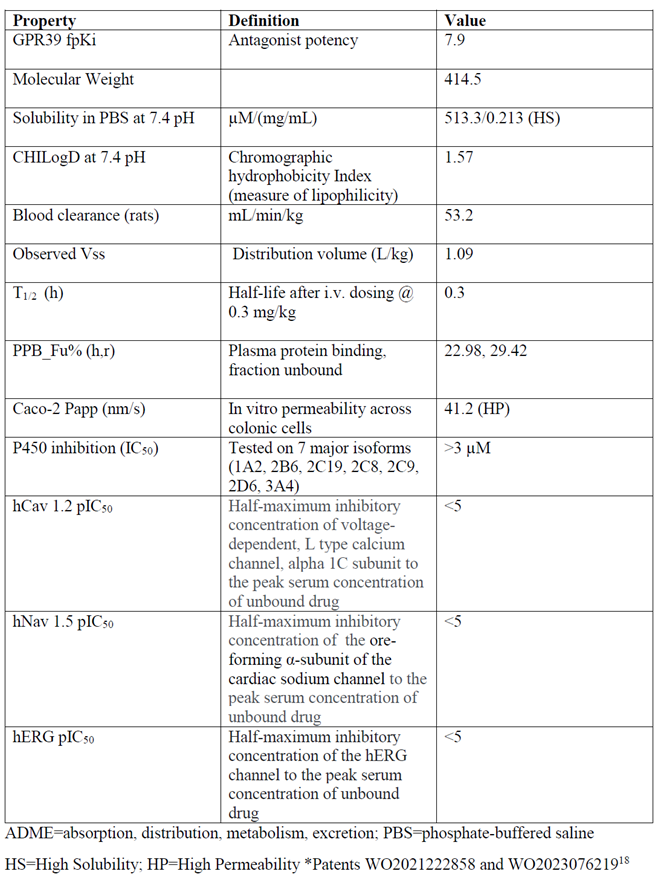

**Supplemental Table 2: Pharmacokinetic Results of VC108 Administered by Intravenous Bolus Injections in Sprague Dawley Rats**

| **Dose (mg/kg/day)** | **Male (n=30)** | | | | **Female (n=30)** | | | |
| --- | --- | --- | --- | --- | --- | --- | --- | --- |
|  | **Mean C_max_**  **(ng/mL)** | | **Mean AUC_last_**  **(ng∙h /mL)** | | **Mean C_max_**  **(ng/mL)** | | **Mean AUC_last_**  **(ng∙h /mL)** | |
|  | **Day 1** | **Day 14** | **Day 1** | **Day 14** | **Day 1** | **Day 14** | **Day 1** | **Day 14** |
| **4 (n=10 M, 10F)** | 3460 | 3990 | 1440 | 2010 | 5460 | 5520 | 5970 | 7030 |
| **15 (n=10 M, 10 F)** | 11100 | 13700 | 4660 | 6520 | 19500 | 23100 | 30500 | 34400 |
| **40 (n=10 M, 10 F)** | 34900 | 42800 | 15100 | 22400 | 55900 | 52700 | 102000 | 135000 |

Of note, these doses of VC108 were markedly higher than those used in the current study.

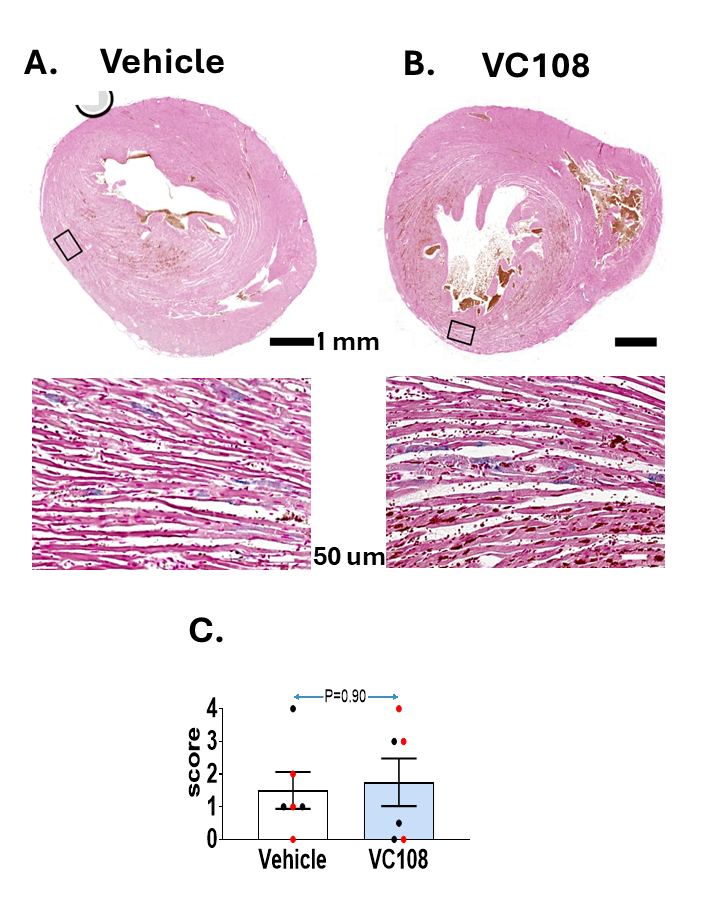
***Supplemental Figure 1:*** *Staining for iron (blue) in Vehicle (A) and VC108 (B) treated animals with aggregate results from 12 animals (C). No difference in iron staining (a marker of ferroptosis) was seen between vehicle and VC108 treated animals.* *Males are depicted in black and females in red.*

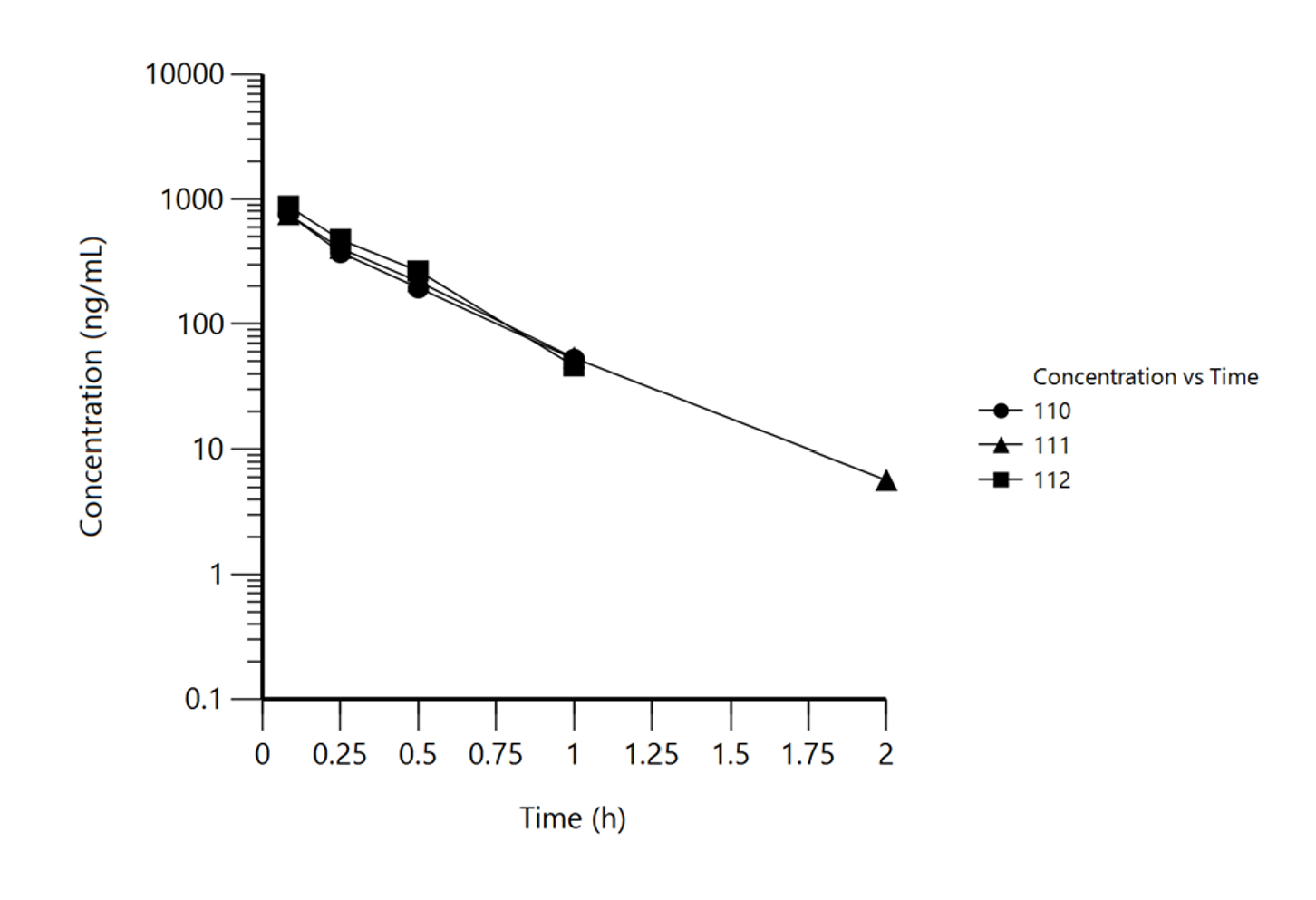
**Supplemental Figure 3**: Time versus Plasma Concentration of VC108 (logarithmic scale) in male rats

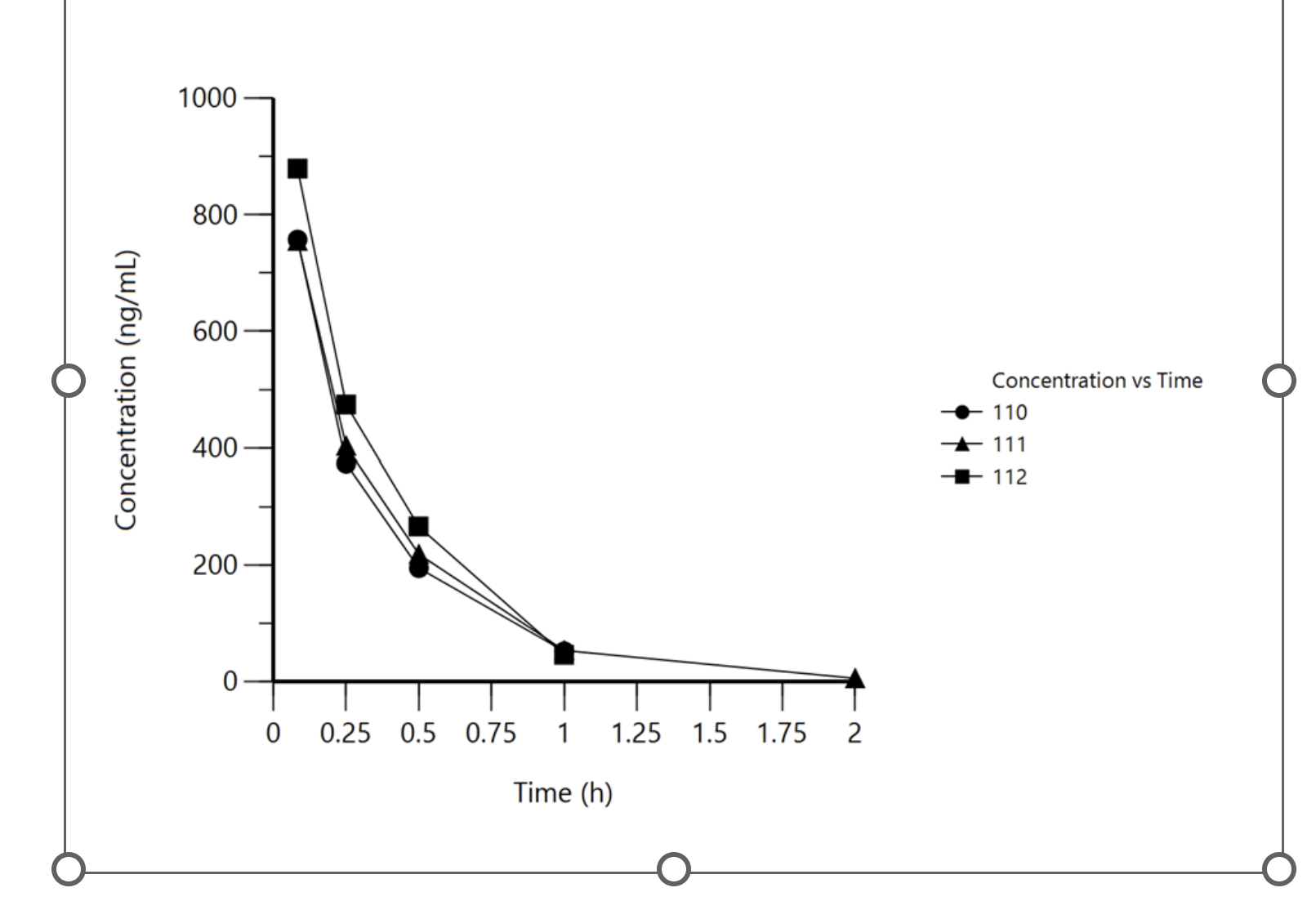
**Supplemental Figure 2**: Time versus Plasma Concentration of VC108 (linear scale) in male rats
